# Pathogenetic Perspective of Missense Mutations of ORF3a Protein of SARS-CoV2

**DOI:** 10.1101/2020.08.04.236653

**Authors:** Sk. Sarif Hassan, Diksha Attrish, Shinjini Ghosh, Pabitra Pal Choudhury, Bidyut Roy

## Abstract

One of the most important proteins for COVID-19 pathogenesis in SARS-CoV2 is the ORF3a protein which is the largest accessory protein among others accessory proteins coded by coronavirus genome. The major roles of the protein include virulence, infectivity, ion channel activity, morphogenesis and virus release. The coronavirus, SARS-CoV2 is continuously evolving naturally and thereby the encoded proteins are also mutating rapidly. Therefore, critical study of mutations in ORF3a is certainty important from the pathogenetic perspective. Here, a sum of 175 various non-synonymous mutations in the ORF3a protein of SARS-CoV2 are identified and their corresponding effects in structural stability and functions of the protein ORF3a are studied. Broadly three different classes of mutations, such as neutral, disease and mixed (neutral and disease) type mutations were observed. Consecutive mutations in some ORF3a proteins are established based on timeline of detection of mutations. Considering the amino acid compositions over the ORF3a primary protein sequences, twenty clusters are detected based on K-means clustering method. Our findings on 175 novel mutations of ORF3a proteins will extend our knowledge of ORF3a, a vital accessory protein in SARS-CoV2, which would assist to enlighten on the pathogenicity of this life-threatening COVID-19.

## 1. Introduction

. Severe Acute Respiratory Syndrome (SARS-CoV) emerged in 2002 infecting about 8000 people with a 10% mortality rate [1, 2]. Similarly, Middle East Respiratory Syndrome Coronavirus (MERS-CoV) emerged in 2012 with 2300 cases and a 35% mortality rate [3]. However, since the December 2019, another outbreak caused by a novel severe acute respiratory syndrome coronavirus 2 (SARS-CoV2) rapidly became a pandemic with the highest mortality rate of 3.4% within just 7 months; urging the World Health Organization to declare it as a Public Health Emergency of International Concern [4, 5, 6, 7]. It was found that SARS-CoV and SARS-CoV2 bear 79% of sequence identity [8, 9]. Similar to SARS-CoV, the ORF3a gene in SARS-CoV2 lies between the spike and envelope gene in virus genome [10]. Both the ORF3a protein of SARS-CoV and SARS-CoV2 contain a conserved cysteine residue which helps in protein-protein interaction [11, 12]. The RNA genome of SARS-CoV2 is about 30 kb in length and codes for 4 structural proteins, 16 non-structural proteins, and 6/7 accessory proteins [13, 14, 15, 16]. The structural proteins are known as Spike protein (S), Nucleocapsid protein (N), Membrane protein (M) and Envelope protein (E) [17].

Among the accessory proteins, our study is based on ORF3a, the largest accessory protein, and a unique membrane protein consisting of three transmembrane domains [18, 19]. SARS-CoV2 ORF3a is a 275 amino acid transmembrane protein that holds an N-terminal, three transmembrane helices followed by a cytosolic domain with multiple *β*-strands [20]. Functionally ORF3a proteins is divided into six domains [21]. Domain I contain N terminus signal peptide involved in subcellular localization of ORF3a protein [19]. Domain II contains a TRAF-3 binding motif (36-40 aa) through which it activates the NF-kB and NLRP3 inflammasome by promoting TNF receptor-associated factor 3 (TRAF3)-mediated ubiquitination of apoptosis-associated speck-like protein containing a caspase recruitment domain (ASC) [21]. Domain III (93-133) is important for ion channel activity and has a Cysteine-rich domain which is associated with homodimerization of ORF3a protein which is very similar to SARS-CoV cysteine rich domain responsible for tetramerization (81-160) [22, 23]. Domain IV has a caveolin binding motif (141-149) which regulates viral uptake and trafficking of protein to the plasma membrane or intracellular membranes [24]. Domain V contains a tyrosine-based sorting motif *Y XXϕ* (160-163) which is responsible for Golgi to plasma membrane transport which in SARS-CoV is responsible for the surface expression [25]. Domain VI has an SGD motif (171-173) [23]. ORF3a has pro-apoptotic activity and membrane association is required for this activity. SARS-CoV2 ORF3a has relatively weaker proapoptotic activity and this property is probably contributing to asymptomatic infection and thus causing rapid transmission of the virus [26]. Therefore, ORF3a may become an important therapeutic target, and thus studying mutations in the ORF3a protein sequence becomes an important area in control of virus infection.

**Figure 1:**
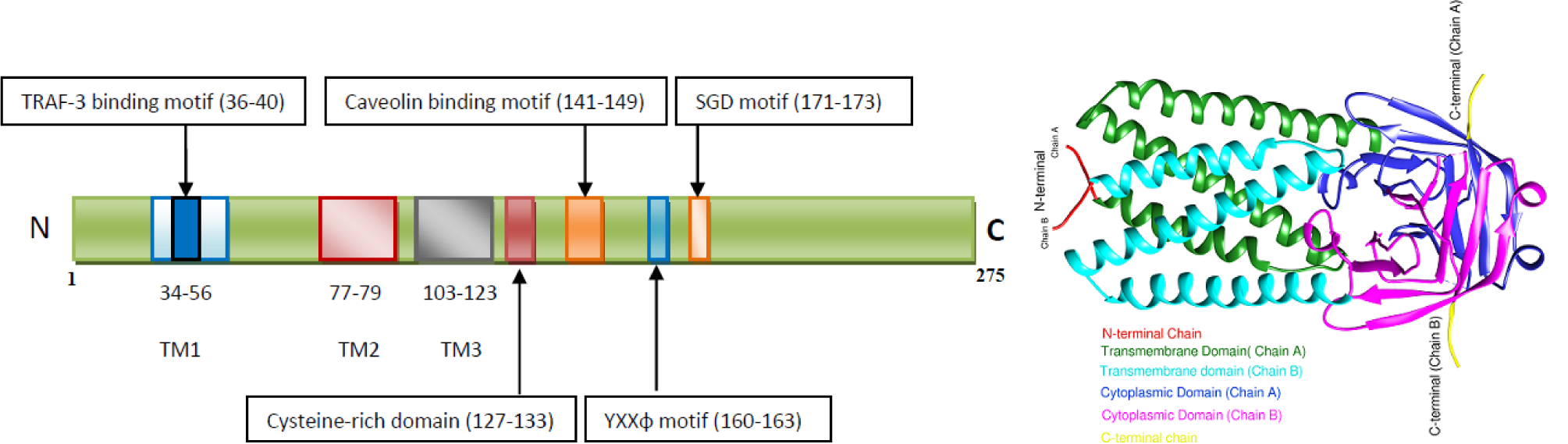
Schematic view of the domains in Primary and Tertiary structures of SARS-CoV2 ORF3a protein.

In our present study, we found about 175 non-synonymous mutations in the ORF3a protein sequence. Among them, 32 are already reported previously [27, 23]. So, we accounted 143 new mutations along with the already existing ones. Mutations in the domain III alters the NF-kB activation and NLRP3 inflammasome. Mutations in domain V were linked to the aggregation of the 3a protein in the Golgi apparatus [28]. Apart from these residues, mutations in 230(insertion of F), W131C, R134L, T151I, N152S and D155Y regions may contribute to a greater significance as they are poised to form a network of hydrophobic, polar and electrostatic interactions which mediate dimerization and tetramerization respectively [29]. To account for mutations of the ORF3a proteins of SARS-CoV2, we collected the SARS-CoV2 genome data from NCBI virus database, identified the mutations, predicted the effect of mutations based on chemical and structural properties. In addition, using the Meta-SNP and I-MUTANT web-servers, effect of the mutations in functions and structures are predicted [30, 31, 32]. We also performed K-means clustering of the distinct variants ORF3a proteins (available as on 27th July, 2020) in order to form twenty disjoint clusters based on the amino acid compositions embedded in the proteins [33, 34]. In addition, Shannon entropy is employed to determine amount of disorderliness of the amino acids over the ORF3a proteins which amplify the wide distinct variations of ORF3a in the USA [35].

## 2. Data and Methods

This present study is based on available genome data of SARS-CoV2 from the NCBI virus database (https://www.ncbi.nlm.nih. Here we discuss about data followed by methods which are employed in this study.

### 2.1. Data

As on date 27th July, 2020, there were 7194 complete genomes of SARS-CoV2 available in the NCBI database and accordingly each genome contains one of the accessory proteins ORF3a and among them only 296 sequences are found to be distinct from each other. The amino acid sequences of ORF3a were exported in fasta format using file operations through Matlab [36]. In this present study, we only concentrate on 296 ORF3a proteins which are listed in the Table-1 and Table-2. Note that, among these 296 sequences, three ORF3a proteins QKO00487 (India: Ahmedabad), QLA10225 (India: Vadodara) and QLA10069 (India: Surat) had the length 241, 253 and 257 respectively and were found to be truncated due to nonsense mutation at 242, 254 and 258 amino acid positions respectively. It is also note worthy that some (13.51%) of 296 ORF3a amino acid sequences contain ambiguous amino acids such as *X, B* and *Z* and so on. In order to find mutations, we hereby consider the reference ORF3a protein as the ORF3a sequence (YP_009724391.1) of the SARS-CoV2 genome (NC_045512) from China: Wuhan [37].

**Table 1:**
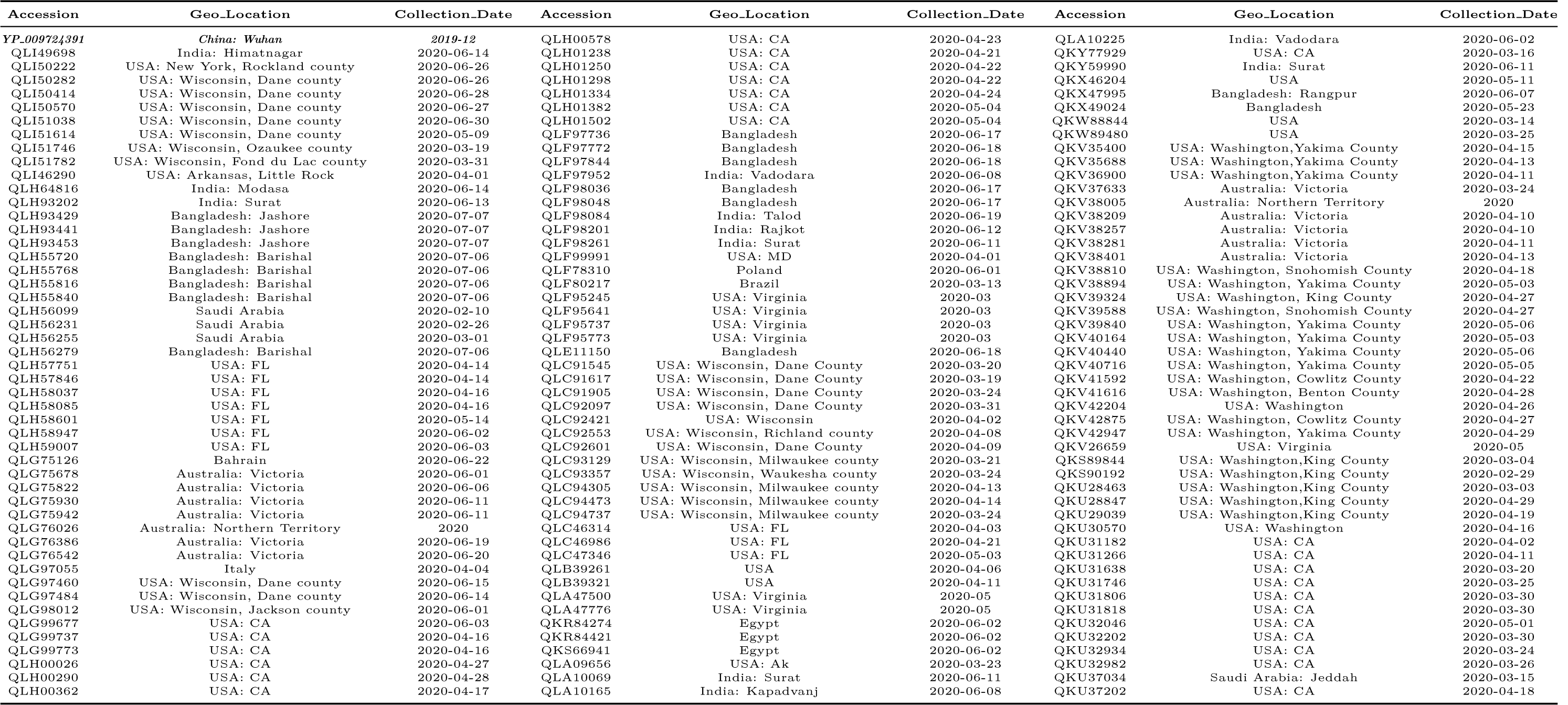
List of accessions of the ORF3a protein, geo-location and respective data collection date.

**Table 2:**
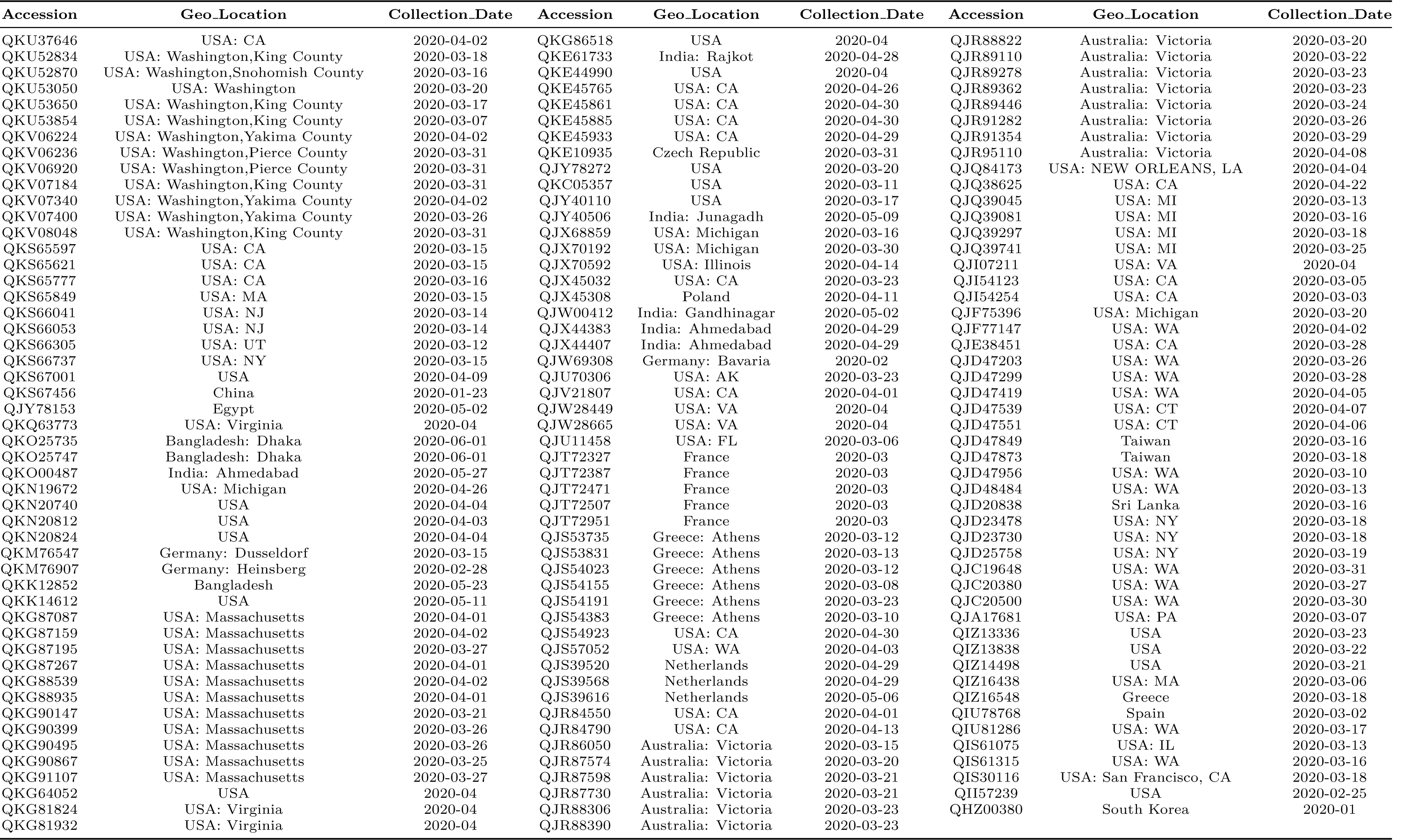
List of accessions of the ORF3a protein, geo-location and respective data collection date.

## 2.2. Methods

Here in a nutshell, we present the methods used in this study as follows.

### 2.2.1. Frequency Probability of Amino Acids

A protein sequence of ORF3a is composed of twenty different amino acids with various frequencies. The probability of occurrence of each amino acid *A*_*i*_ is determined by the formula 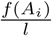 where *f* (*A*_*i*_) denotes the frequency of occurrence of the amino acid *A*_*i*_ in the primary sequence ORF3a and *l* stands as the length of ORF3a protein [38]. Hence for each of the 296 ORF3a proteins, a twenty dimensional vector considering the frequency probability of twenty amino acids can be obtained. Based on these frequency probability vectors, a classification is performed using clustering technique.

### 2.2.2. K-means Clustering Algorithm

Clustering is one of the most widely used methods in vector-data analysis to develop an intuitive idea about closeness of data based on the structured feature vectors. By clustering we find homogeneous subclasses within the data such that data points in each cluster are as similar as possible according to a similarity measure such as euclidean-based distance. One of the most commonly used simple clustering techniques is the *K-means clustering* [33, 34].

**Algorithm:** K-means algorithm is an iterative algorithm that tries to form equivalence classes from the feature vectors into K (pre-defined) clusters where each data point belongs to only one cluster [33].

- Assign the number of desired clusters *(K*) (in the present study, *K* = 20).
- Finding centroids by first shuffling the dataset and then randomly selecting *K* data points for the centroids without replacement.
- Keep iterating until there is no change to the centroids.
- Find the sum of the squared distance between data points and all centroids.
- Assign each data point to the closest cluster (centroid).
- Compute the centroids for the clusters by taking the average of the all data points that belong to each cluster.

In this study we did clustering using Matlab by customizing the value of K and inputting the frequency of amino acid compositions over the ORF3a proteins.

### 2.2.3. Amino Acid Conservation Shannon Entropy

How conserved/disordered the amino acids are, over ORF3a protein is addressed by the information theoretic measure known as ‘*Shannon entropy*(SE)’ which we deploy here to find out conservation entropy of each ORF3a protein. For each ORF3a protein, Shannon entropy of amino acid conservation over the amino acid sequence of ORF3a protein is computed 85 using the following formula [39]:

For a given amino acid sequence of ORF3a protein of length *l*, the conservation of amino acids is calculated as follows:

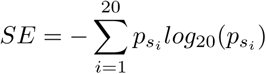

where 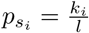; *k*_*i*_ represents the number of occurrences of an amino acid *s*_*i*_ in the given sequence.

In this study, SE describes the wide variety of 296 distinct ORF3a proteins collected from various countries across the world.

## 3. Results

All mutations, compared to Chinese Wuhan sequence, over the set of distinct ORF3a proteins are detected and consequently they have been classified based on their predicted effect as disease/neutral in important functions of ORF3a protein (Table 9). Also, some important known domains are identified for the observed mutations and accordingly the predicted effect of mutations in protein functions have been discussed. Further, consecutive mutations observed in ORF3a proteins according to the timelines of detection of various mutations for a subgroup of ORF3a proteins located in Australia, Bangladesh, India, USA and so on is derived (Fig.7-11). Using a web-server (i − MUTANT: http://gpcr2.biocomp.unibo.it/cgi/predictors/I − Mutant3.0/I − Mutant3.0.cgi) stability of ORF3a protein structures were predicted upon various mutations. At last, twenty clusters are formed using K-means clustering method based on frequency probability of amino acids of 296 ORF3a proteins. The wide variations of 296 ORF3a proteins are finally supported by the Shannon entropy (SE) and remarkably we found the most widest varieties of ORF3a proteins in virus detected in the USA.

### 3.1. Mutations over the ORF3a protein of SARS-CoV2

Each of the ORF3a amino acid sequences (fasta formatted) are aligned with respect to the ORF3a protein (YP_009724391.1) from China-Wuhan using multiple sequence alignment tool (NCBI Blastp suite) and found the mutations and their associated positions were detected accordingly [40]. It is noted that a mutation from an amino acid *A*_1_ to *A*_2_ at a position *p* is denoted by *A*_1_*pA*_2_ or *A*_1_(*p*)*A*_2_. The Fig.2 describes various mutations with their respective locations. The mutations are found in the entire ORF3a sequence starting from the amino acid position 7 to 271. It is found that an amino acid at a fixed position mutates to two different amino acids. For examples, at 9th position of the reference ORF3a protein, the amino acid Threonine(T) maps to Isoleucine(I) and Lysine(K) in different ORF3a proteins. At the 18th position Glycine maps to three amino acids Valine, Serine and Cysteine. The amino acid Alanine(A) maps to Valine, Serine, Threonine and Aspartic acid at the 99th position.

**Figure 2:**
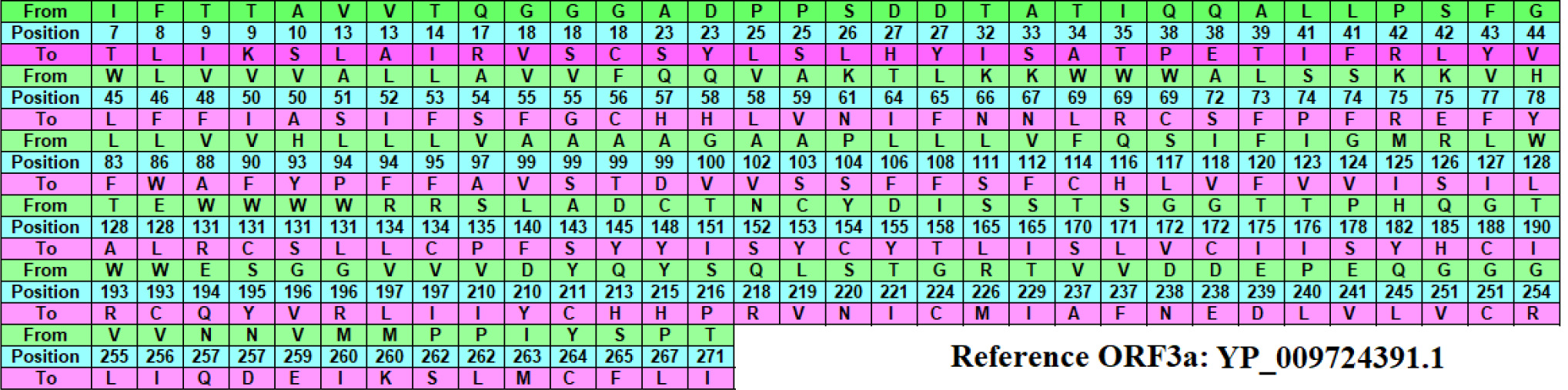
Mutations in the respective position in ORF3a protein sequence compared with reference Wuhan sequence YP_009724391.1. Note: From: existing amino acid in reference sequence; position: amino acid position in the sequence; To: mutated amino acid in studied sequence

Based on observed mutations, it is noticed that amino acids Alanine(A) and Tryptophan(W) are found to be most vulnerable to mutate to various amino acids. It is noted that the mutation of Tryptophan (W) at 131 position are found in the Cystine-rich domain (127-133).

Distinct mutations and its associated mutation of frequency are presented in Table-3. The most frequent mutation over the ORF3a is to be Q57H (Acidity: Neutral(Q) to Basic(weakly)(H)) with frequency 142. A pie chart accounting the frequency distribution of various mutations is shown in Fig.3. In addition to the list of mutations (Fig.2), two deletion and two insertion mutations were found in five different ORF3a proteins at various positions.

**Figure 3:**
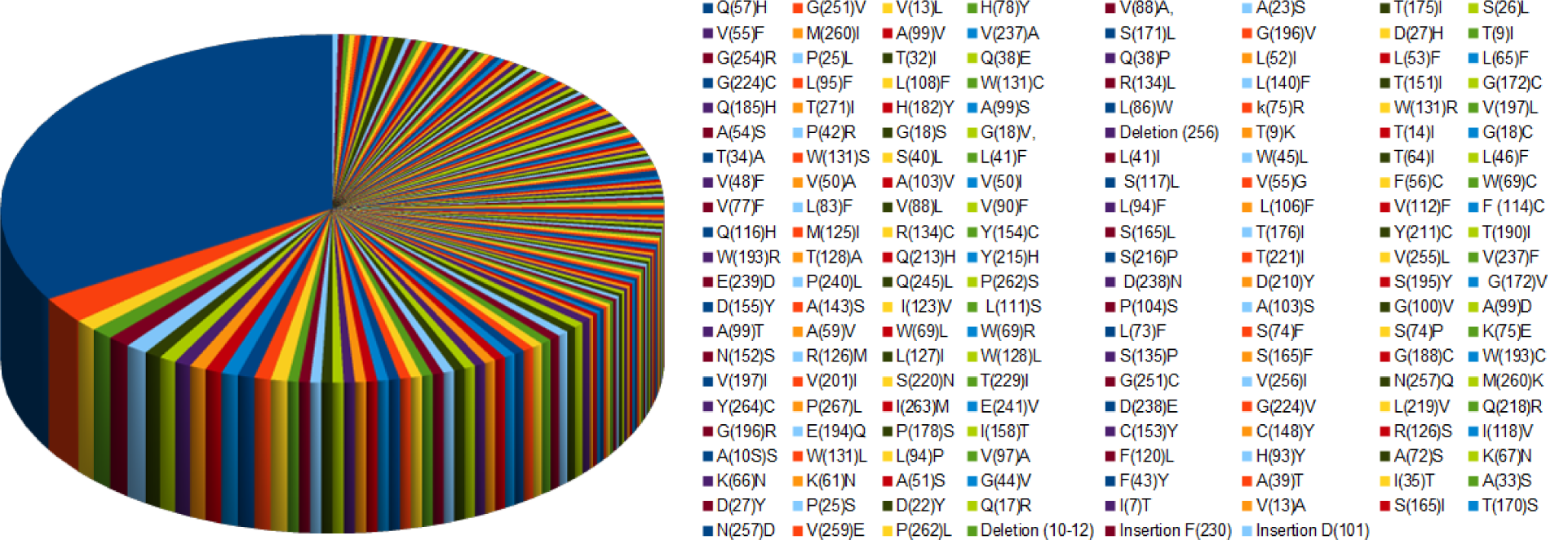
Pie chart of the frequency of distinct mutations.

The details of mutations, in the 256 ORF3a unique proteins from viruses of 256 patients, in specific domain(s) and predicted effects of mutations viz. disease and neutral effects through the web-server Meta-SNP (https://snps.biofold.org/meta-snp/) are presented in the Tables-4,5, 6, 7 & 8. Note that among 296 ORF3a proteins, 40 sequences possess only ambiguous mutations which we have neglected. A snapshot of predicted result (disease causing variant with reliability score 3) of the most frequent mutation Q57H is shown in Fig.4.

**Table 3:**
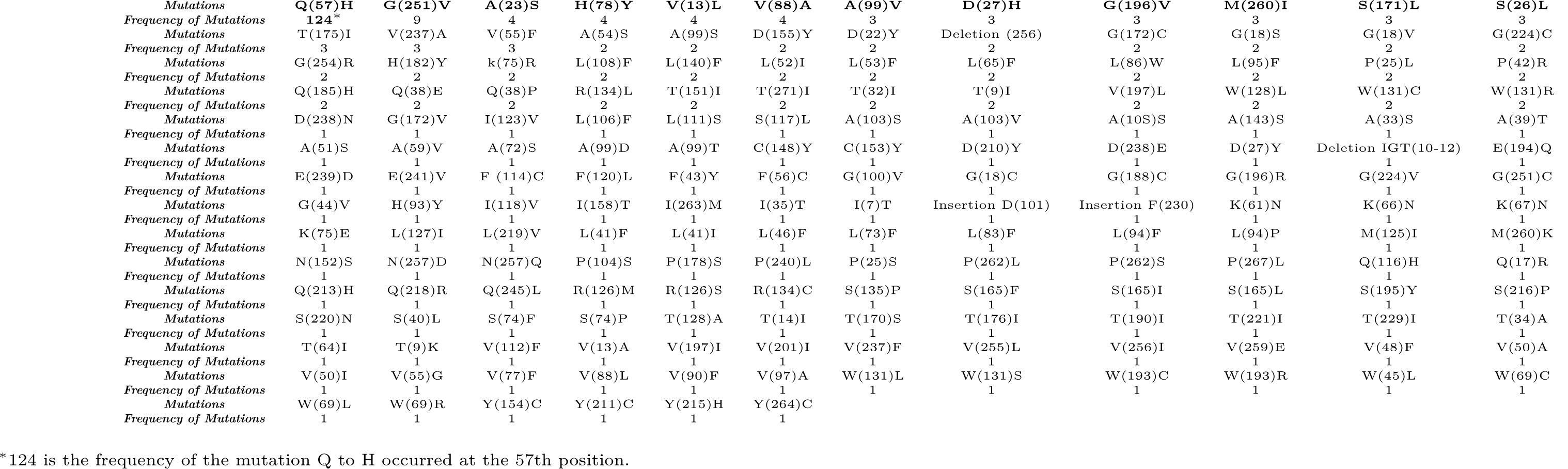
Distinct mutations across the ORF3a proteins and their respective frequency.

**Figure 4:**
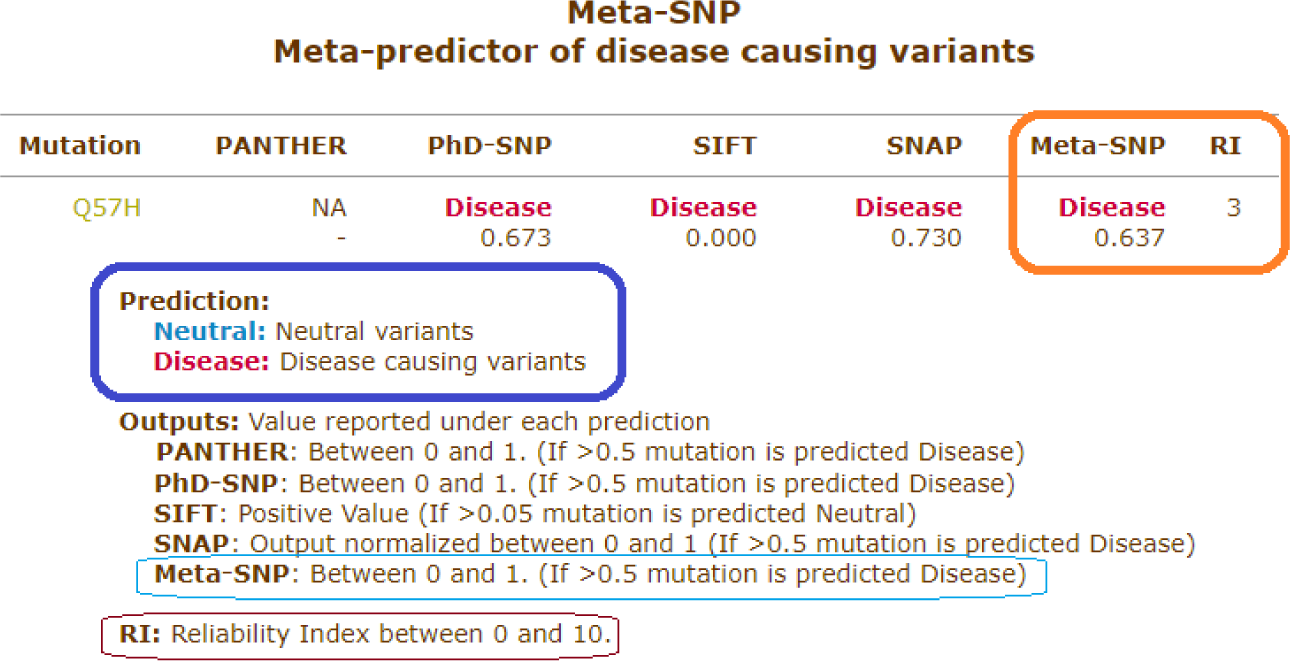
A snapshot of the predicted effect of the frequently occurred mutation Q57H in ORF3a using Meta-SNP web-server.

Based on the predicted type of mutations, all the 256 ORF3a proteins are classified into three classes which are presented in the Table 9. The three classes representing disease, neutral and mixture of disease as well as neutral mutations are constituted of protein IDs with respective geo-locations.

**Table 4:**
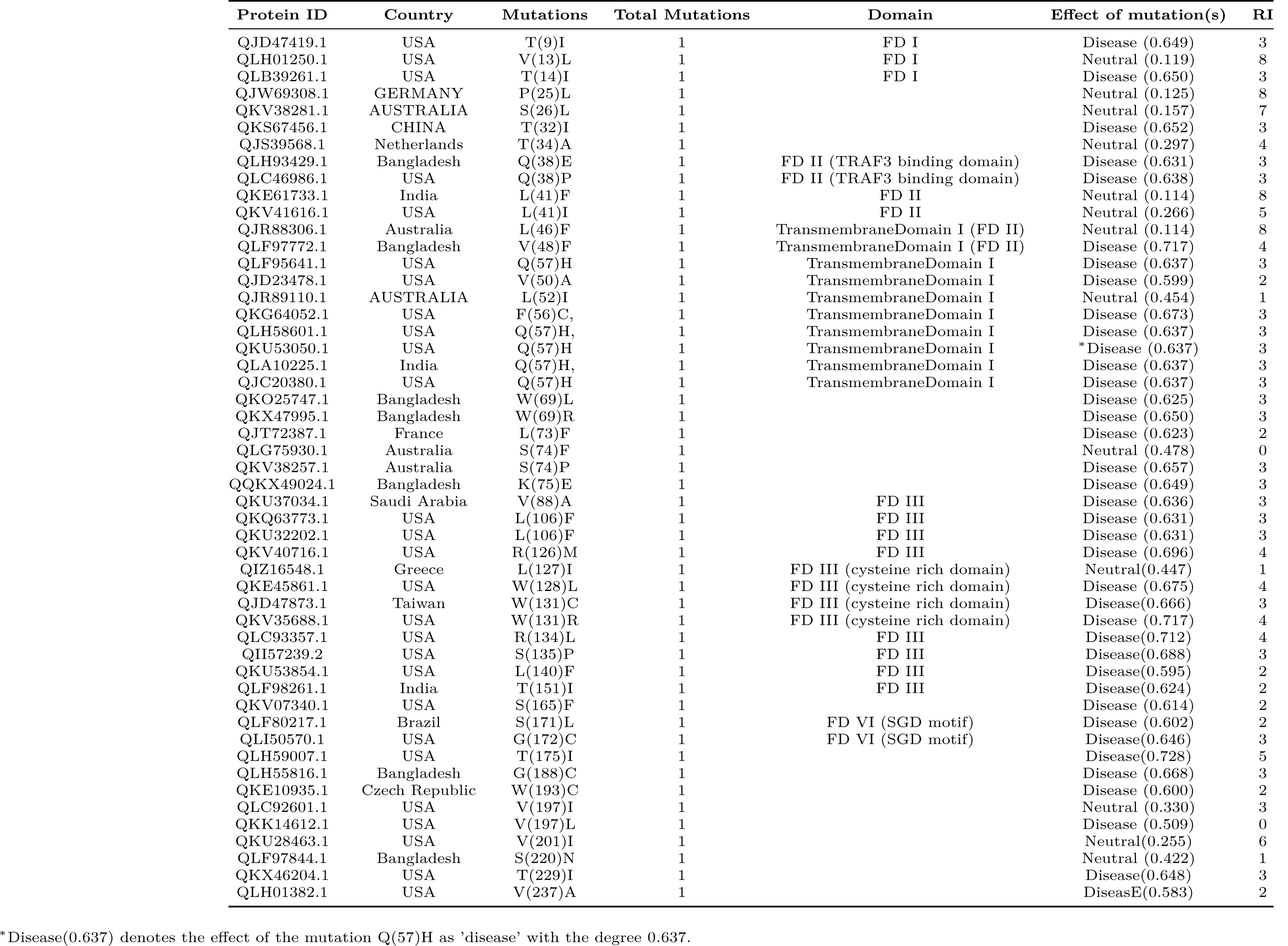
protein IDs and respective mutations, geo-locations, total number of mutations in the protein, domains and predicted effect of the mutations.

**Table 5:**
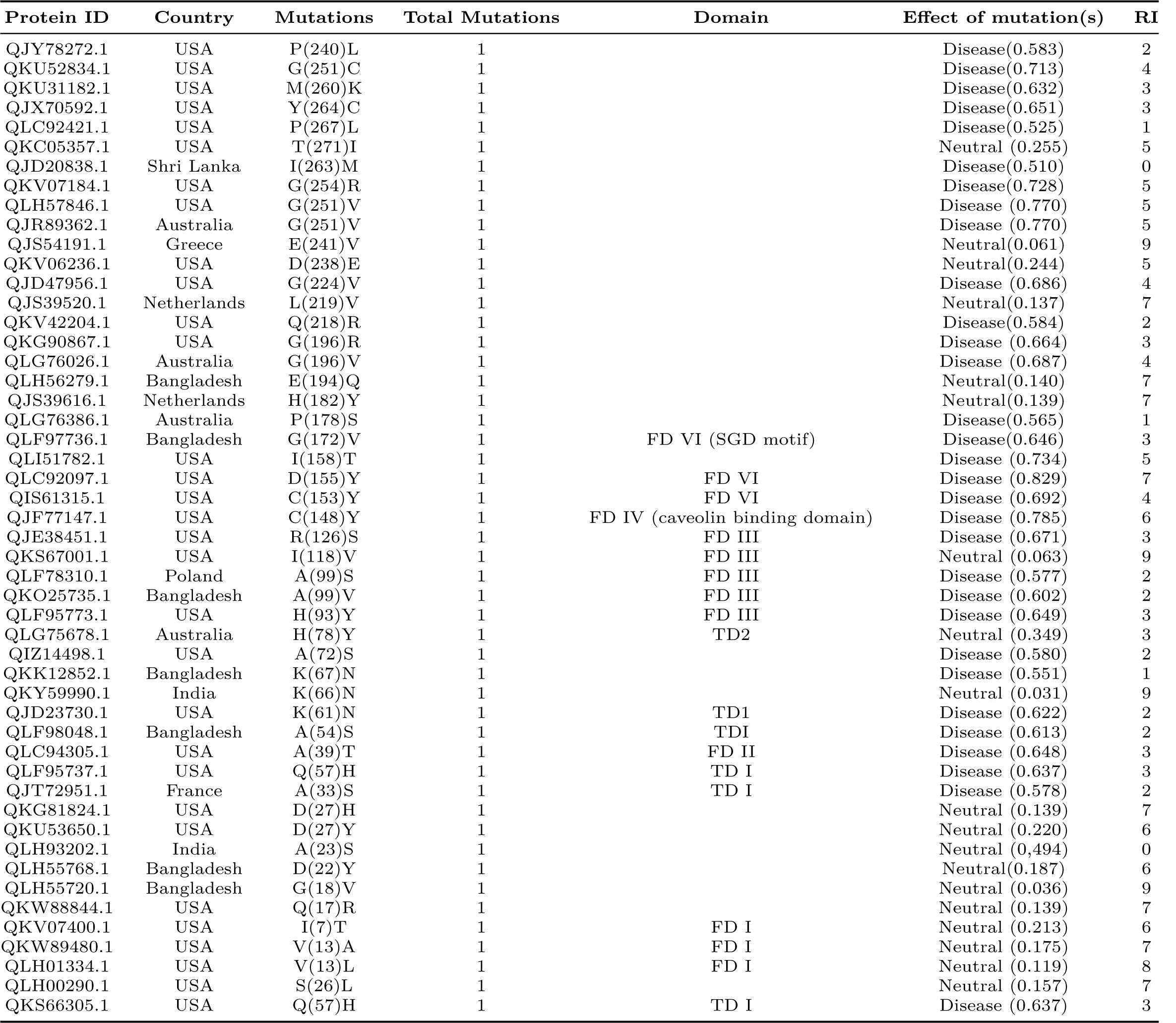
protein IDs and respective mutations, geo-locations, total number of mutations, domain and predicted effect of the mutations.

**Table 6:**
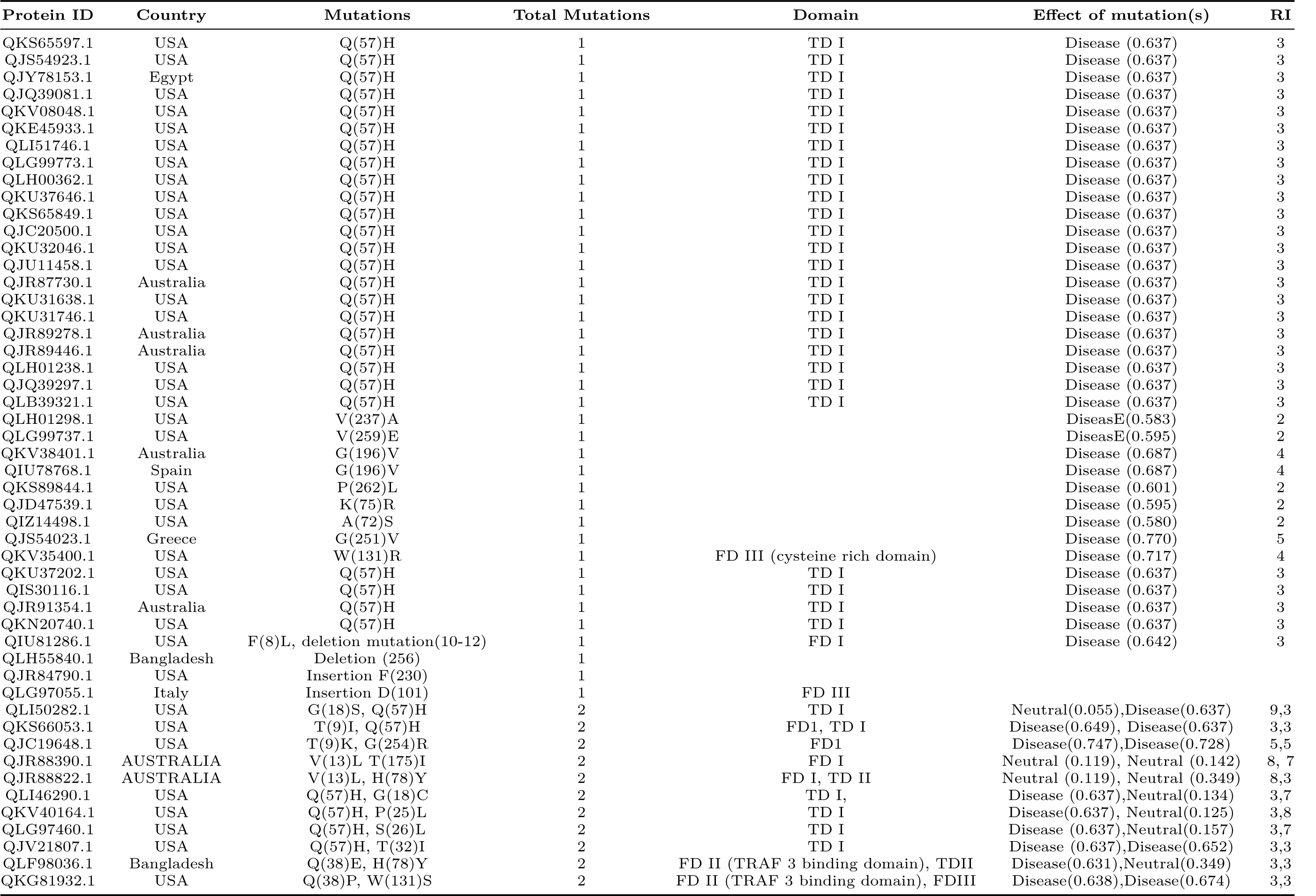
protein IDs and respective mutations, geo-locations, total number of mutations, domain and predicted effect of the mutations.

**Table 7:**
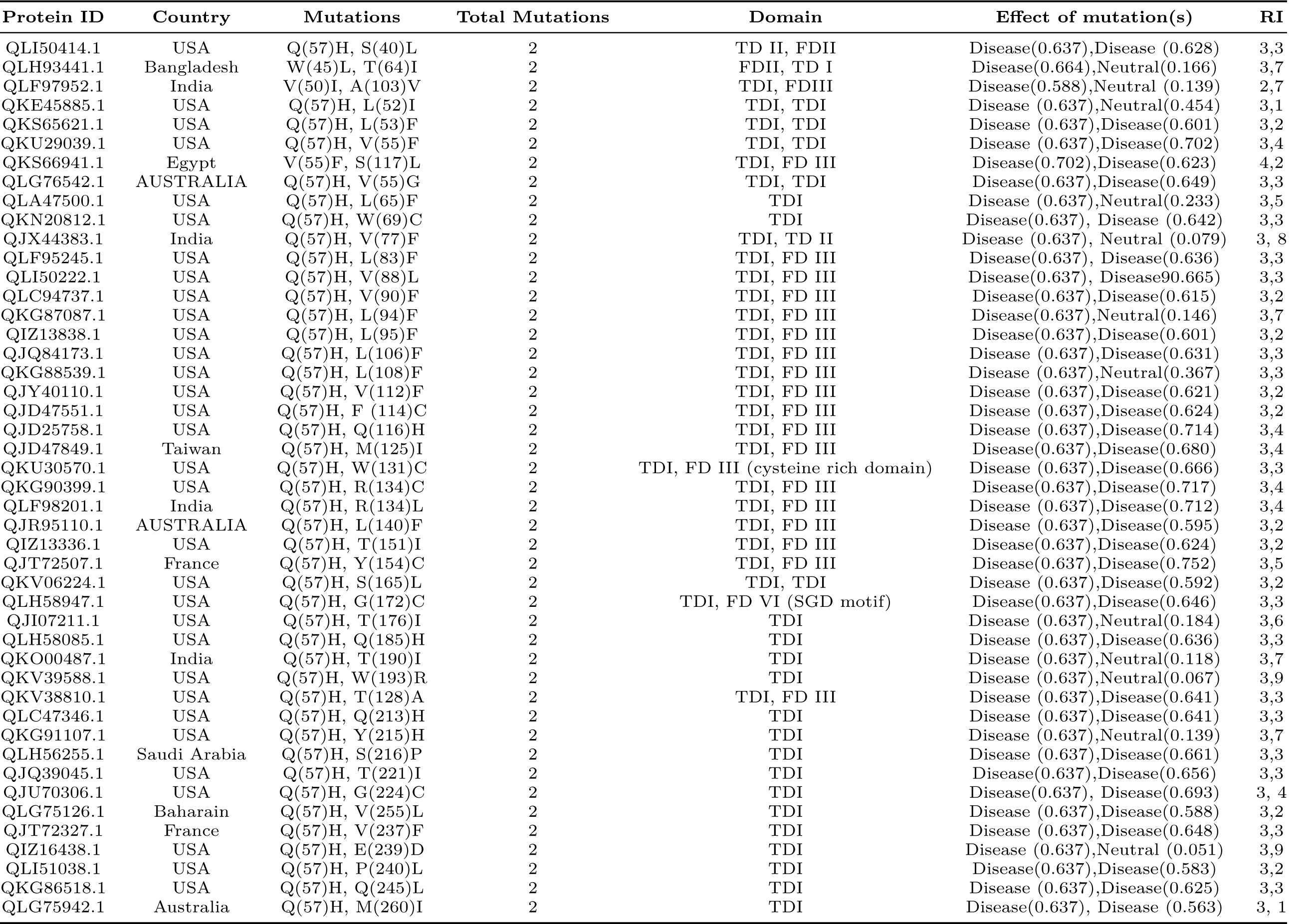
protein IDs and respective mutations, geo-locations, total number of mutations, domain and predicted effect of the mutations.

**Table 8:**
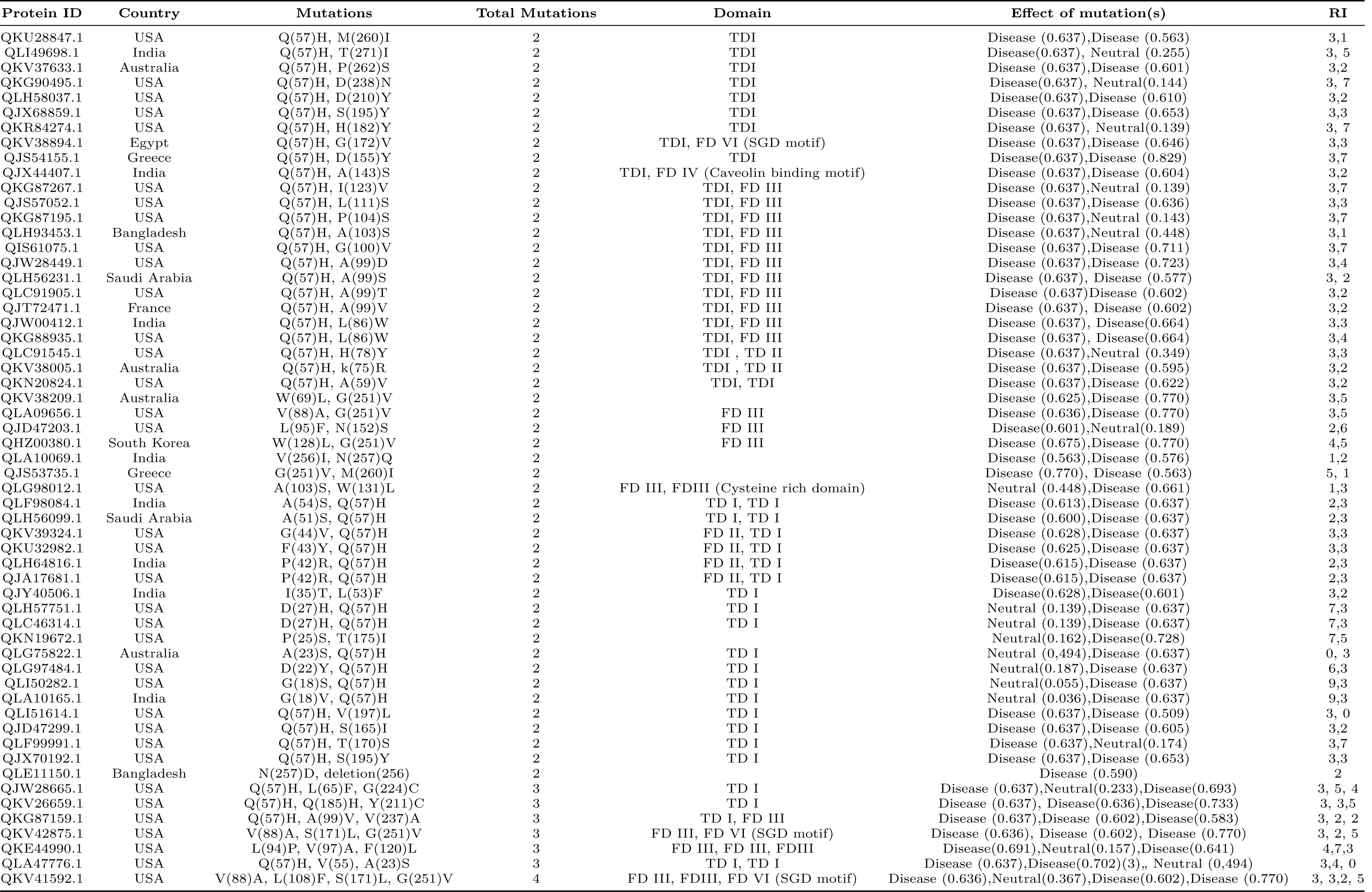
protein IDs and respective mutations, geo-locations, total number of mutations, domain and predicted effect of the mutations.

**Table 9:**
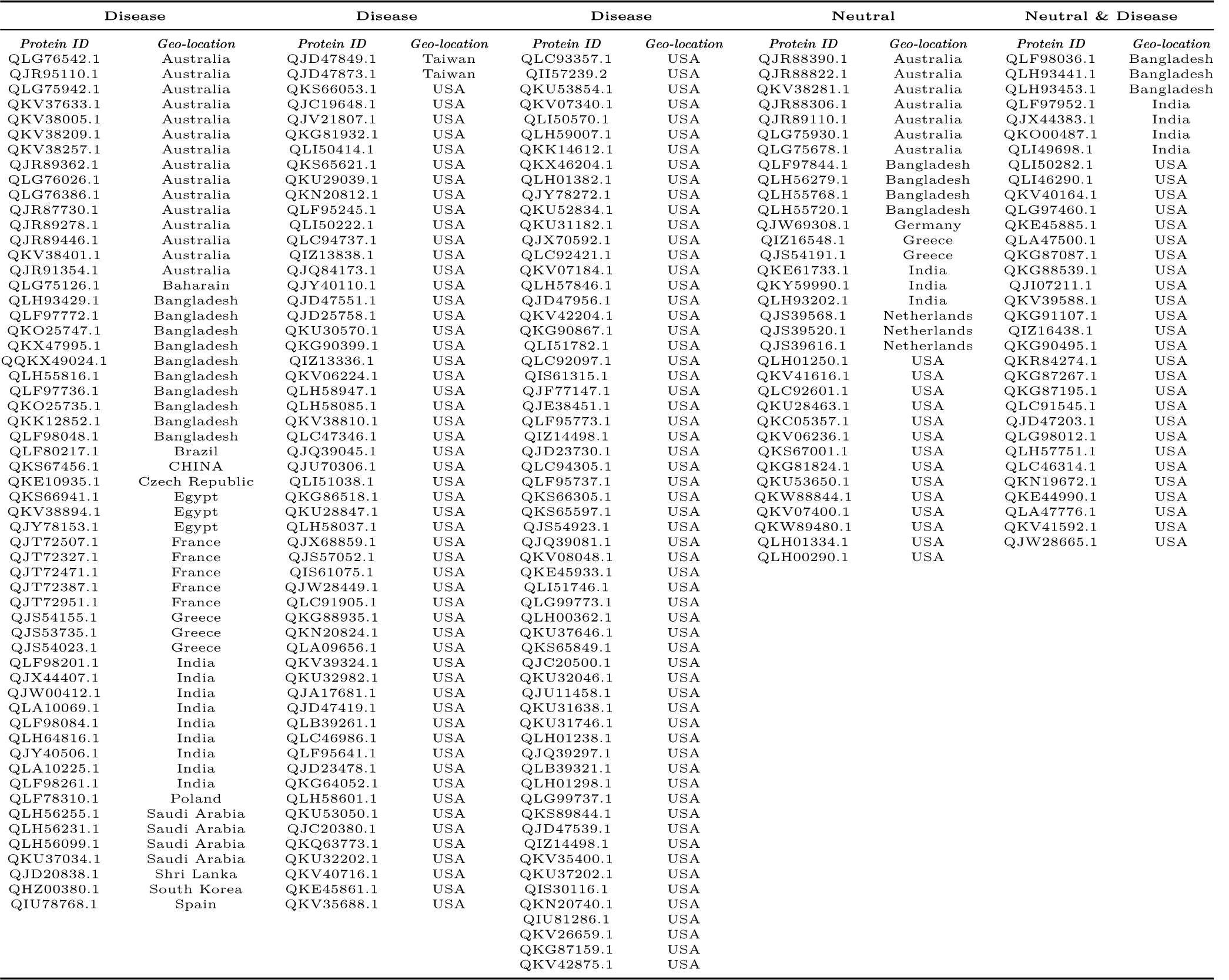
ORF3a proteins possessed disease, neutral and mixed of neutral & disease type of predicted mutations.

Almost 72% of the ORF3a proteins possess disease type of mutations whereas 14% (of which two mutations: 12%, three mutations: 1.5% and four mutations: 0.5%) and 14% of ORF3a proteins possess mixture type (i.e. both disease as well as neutral) and neutral types of mutations respectively (Fig.5).

**Figure 5:**
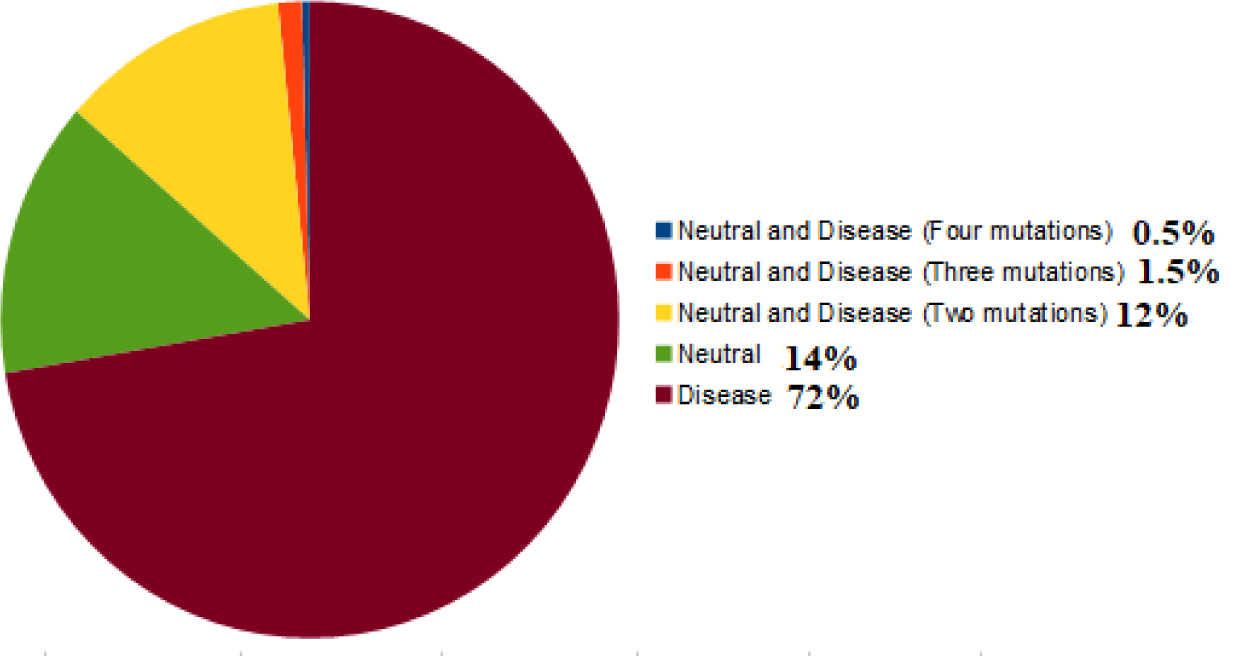
Percentage of disease, neutral and mix (neutral & disease) type of mutations over the ORF3a proteins.

For each of the three types of mutations, we put the frequency and percentage of ORF3a proteins corresponding to each geo-locations as presented in the Table 10.

In USA, all three type of mutations over the ORF3a proteins are found to be dominant in percentage. In a Fig.6, the world maps are marked as per occurrence of three types of mutations in ORF3a variants.

**Figure 6:**
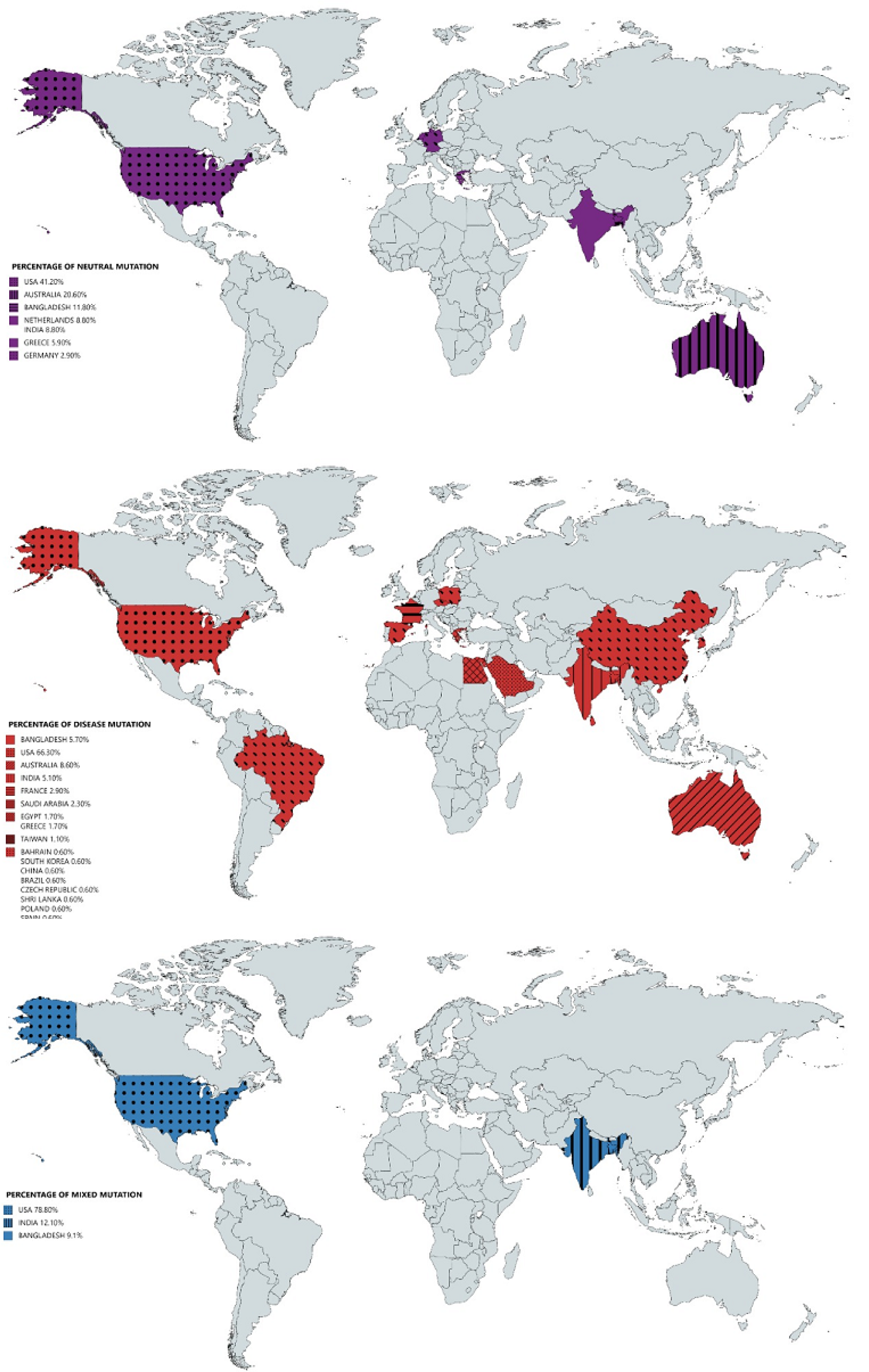
World maps of percentage of occurrence of neutral, disease and mixed type of mutations over the ORF3a proteins.

Frequency of neutral mutation is 41.2% being the highest in the USA, according to prediction it shows that this mutation is neutral but still this mutation is supposed to be contributing to the weaker apoptotic activity of ORF3a and this weaker activity may be responsible for asymptomatic or relatively mildly symptomatic cases thus causing rapid transmission of the virus.

### 3.2. Possible Consecutive mutations over ORF3a proteins

Several ORF3a proteins (Tables 4-8) contain more than one mutations and maximally up to four mutations. It takes time for multiple mutations in a given ORF3a protein and relying on time-line and order occurrence of mutations several flow of consecutive mutations were derived. The predicted effects of these mutations on stability of the tertiary structure of the ORF3a proteins was determined in the flow of consecutive mutations (Table 11).

**Table 10:** Frequency and percentage of ORF3a proteins located at various countries, having three type of mutations.

| Disease |  |  | Neutral |  |  | Neutral & Disease |  |  |
| --- | --- | --- | --- | --- | --- | --- | --- | --- |
| *Geo-location* | *Frequency* | *Percentage* | *Geo-location* | *Frequency* | *Percentage* | *Geo-location* | *Frequency* | *Percentage* |
| USA | 116 | 66.30% | USA | 14 | 41.20% | USA | 26 | 78.80% |
| AUSTRALIA | 15 | 8.60% | AUSTRALIA | 7 | 20.60% | INDIA | 4 | 12.10% |
| BANGLADESH | 10 | 5.70% | BANGLADESH | 4 | 11.80% | BANGLADESH | 3 | 9.10% |
| INDIA | 9 | 5.10% | NETHERLANDS | 3 | 8.80% |  |  |  |
| FRANCE | 5 | 2.90% | INDIA | 3 | 8.80% |  |  |  |
| SAUDI ARABIA | 4 | 2.30% | GREECE | 2 | 5.90% |  |  |  |
| EGYPT | 3 | 1.70% | GERMANY | 1 | 2.90% |  |  |  |
| GREECE | 3 | 1.70% |  |  |  |  |  |  |
| TAIWAN | 2 | 1.10% |  |  |  |  |  |  |
| BAHARAIN | 1 | 0.60% |  |  |  |  |  |  |
| SOUTH KOREA | 1 | 0.60% |  |  |  |  |  |  |
| CHINA | 1 | 0.60% |  |  |  |  |  |  |
| BRAZIL | 1 | 0.60% |  |  |  |  |  |  |
| CZECH REPUBLIC | 1 | 0.60% |  |  |  |  |  |  |
| SHRI LANKA | 1 | 0.60% |  |  |  |  |  |  |
| POLAND | 1 | 0.60% |  |  |  |  |  |  |
| SPAIN | 1 | 0.60% |  |  |  |  |  |  |

**Table 11:**
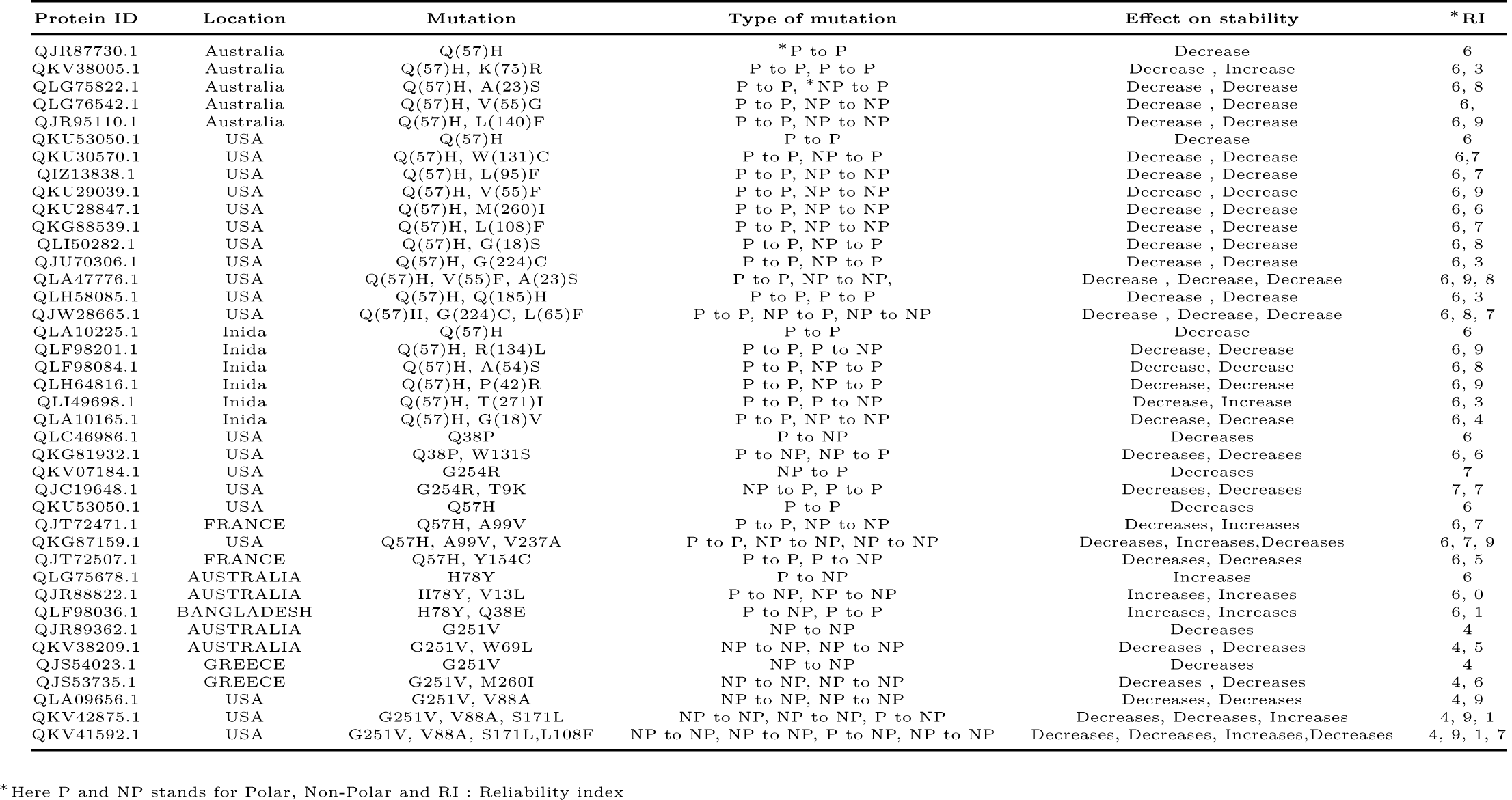
ORF3a proteins with associated mutations and predicted effect in stability of the structures.

| Protein ID | Location | Mutation | Type of mutation | Effect on stability | * RI |
| --- | --- | --- | --- | --- | --- |
| QJR87730.1 | Australia | Q(57)H | * P to P | Decrease | 6 |
| QKV38005.1 | Australia | Q(57)H, K(75)R | P to P, P to P | Decrease , Increase | 6, 3 |
| QLG75822.1 | Australia | Q(57)H, A(23)S | P to P, *NP to P | Decrease , Decrease | 6, 8 |
| QLG76542.1 | Australia | Q(57)H, V(55)G | P to P, NP to NP | Decrease , Decrease | 6, |
| QJR95110.1 | Australia | Q(57)H, L(140)F | P to P, NP to NP | Decrease , Decrease | 6, 9 |
| QKU53050.1 | USA | Q(57)H | P to P | Decrease | 6 |
| QKU30570.1 | USA | Q(57)H, W(131)C | P to P, NP to P | Decrease , Decrease | 6, 7 |
| QIZ13838.1 | USA | Q(57)H, L(95)F | P to P, NP to NP | Decrease , Decrease | 6, 7 |
| QKU29039.1 | USA | Q(57)H, V(55)F | P to P, NP to NP | Decrease , Decrease | 6, 9 |
| QKU28847.1 | USA | Q(57)H, M(260)I | P to P, NP to NP | Decrease , Decrease | 6, 6 |
| QKG88539.1 | USA | Q(57)H, L(108)F | P to P, NP to NP | Decrease , Decrease | 6, 7 |
| QLI50282.1 | USA | Q(57)H, G(18)S | P to P, NP to P | Decrease , Decrease | 6, 8 |
| QJU70306.1 | USA | Q(57)H, G(224)C | P to P, NP to P | Decrease , Decrease | 6, 3 |
| QLA47776.1 | USA | Q(57)H, V(55)F, A(23)S | P to P, NP to NP, | Decrease , Decrease, Decrease | 6, 9, 8 |
| QLH58085.1 | USA | Q(57)H, Q(185)H | P to P, P to P | Decrease , Decrease | 6, 3 |
| QJW28665.1 | USA | Q(57)H, G(224)C, L(65)F | P to P, NP to P, NP to NP | Decrease , Decrease, Decrease | 6, 8, 7 |
| QLA10225.1 | Inida | Q(57)H | P to P | Decrease | 6 |
| QLF98201.1 | Inida | Q(57)H, R(134)L | P to P, P to NP | Decrease, Decrease | 6, 9 |
| QLF98084.1 | Inida | Q(57)H, A(54)S | P to P, NP to P | Decrease, Decrease | 6, 8 |
| QLH64816.1 | Inida | Q(57)H, P(42)R | P to P, NP to P | Decrease, Decrease | 6, 9 |
| QLI49698.1 | Inida | Q(57)H, T(271)I | P to P, P to NP | Decrease, Increase | 6, 3 |
| QLA10165.1 | Inida | Q(57)H, G(18)V | P to P, NP to NP | Decrease, Decrease | 6, 4 |
| QLC46986.1 | USA | Q38P | P to NP | Decreases | 6 |
| QKG81932.1 | USA | Q38P, W131S | P to NP, NP to P | Decreases, Decreases | 6, 6 |
| QKV07184.1 | USA | G254R | NP to P | Decreases | 7 |
| QJC19648.1 | USA | G254R, T9K | NP to P, P to P | Decreases, Decreases | 7, 7 |
| QKU53050.1 | USA | Q57H | P to P | Decreases | 6 |
| QJT72471.1 | FRANCE | Q57H, A99V | P to P, NP to NP | Decreases, Increases | 6, 7 |
| QKG87159.1 | USA | Q57H, A99V, V237A | P to P, NP to NP, NP to NP | Decreases, Increases, Decreases | 6, 7, 9 |
| QJT72507.1 | FRANCE | Q57H, Y154C | P to P, P to NP | Decreases, Decreases | 6, 5 |
| QLG75678.1 | AUSTRALIA | H78Y | P to NP | Increases | 6 |
| QJR88822.1 | AUSTRALIA | H78Y, V13L | P to NP, NP to NP | Increases, Increases | 6, 0 |
| QLF98036.1 | BANGLADESH | H78Y, Q38E | P to NP, P to P | Increases, Increases | 6, 1 |
| QJR89362.1 | AUSTRALIA | G251V | NP to NP | Decreases | 4 |
| QKV38209.1 | AUSTRALIA | G251V, W69L | NP to NP, NP to NP | Decreases , Decreases | 4, 5 |
| QJS54023.1 | GREECE | G251V | NP to NP | Decreases | 4 |
| QJS53735.1 | GREECE | G251V, M260I | NP to NP, NP to NP | Decreases , Decreases | 4, 6 |
| QLA09656.1 | USA | G251V, V88A | NP to NP, NP to NP | Decreases, Decreases | 4, 9 |
| QKV42875.1 | USA | G251V, V88A, S171L | NP to NP, NP to NP, P to NP | Decreases, Decreases, Increases | 4, 9, 1 |
| QKV41592.1 | USA | G251V, V88A, S171L, L108F | NP to NP, NP to NP, P to NP, NP to NP | Decreases, Decreases, Increases, Decreases | 4, 9, 1, 7 |

\*Here P and NP stands for Polar, Non-Polar and RI : Reliability index

#### Flow of consecutive mutation-I

In the Australian region, it can be observed that the first mutation may have occurred in sequence QJR87730.1 with respect to the Wuhan sequence (YP_009724391.1) from Q to H at 57th position which is a disease type mutation and also this mutation is having the highest frequency which may indicate that it has an important role to play in infectivity part of the virus. As we move along the flow, six ORF3a sequences were considered based on the consecutive time scale of detection that was found to have 2nd mutation on the background of initial Q57H mutation with reference to Wuhan sequence (YP_009724391.1) (Fig.7).

**Figure 7:**
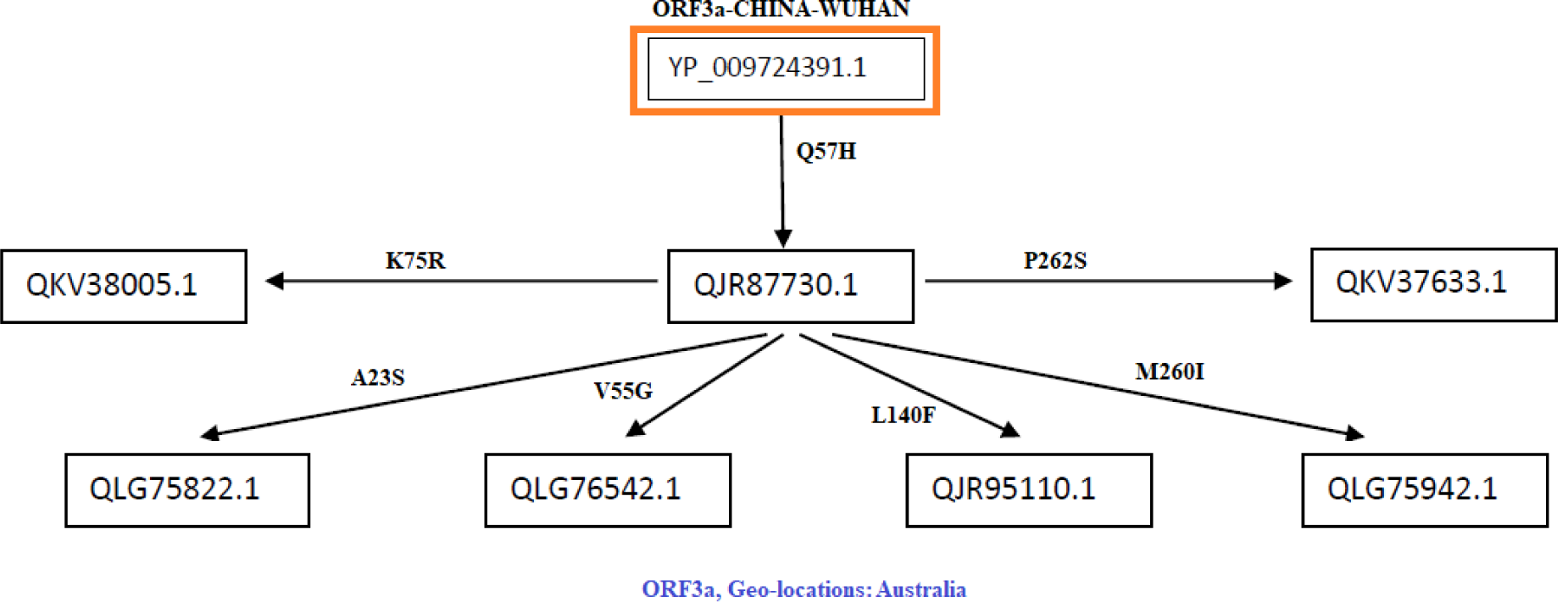
Flow of mutations in Australian ORF3a proteins.

In this flow of mutation, six ORF3a proteins possess various mutations as follows:

- In QKV38005.1, there is a mutation K75R which was found to be a diseased type. We have to consider disease type mutation which may change the function of the protein.
- In QLG75822.1, there is a mutation A23S which was found to be a neutral type with no polarity change. So this is a synonymous mutation from the functionality perspective.
- In QLG76542.1, there is a mutation V55G which was found to be a diseased type, and and hydrophobicity changed to hydrophilicity. This indicates that there may be a functional importance of this mutation.
- In QJR95110.1, there is a mutation L140F which was found to be a diseased type with no polarity change. Since no polarity change is observed the type of amino acid remains same but the mutation effect becomes harmful for the host.
- In QLG75942.1, there is a mutation at M260I that was found to be a diseased type with no polarity change. This mutation may increase the virus virulence.
- In QKV37633.1, there is a mutation at P262S which was found to be a diseased type, and polarity changed from hydrophobic to hydrophilic. Consequently, it may account for change in structure of the protein.

#### Flow of consecutive mutation-II

The most frequent mutation Q57H occurred in the ORF3a protein QKU53050.1. In this network flow (Fig.8) there are other nine sequences which are considered based on the succeeding time scale that was found to have 2nd level mutations along with Q57H.

**Figure 8:**
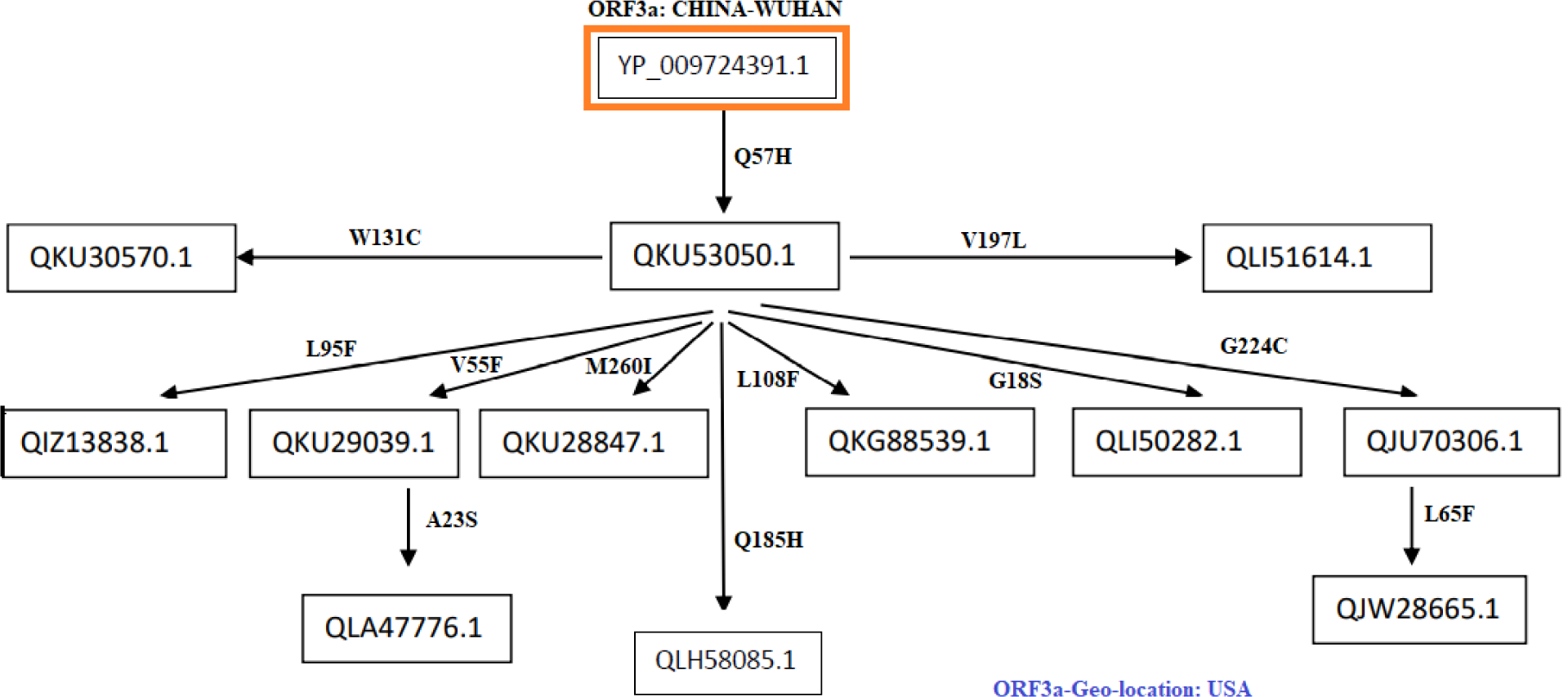
Flow of mutations in ORF3a proteins from the USA.

- The ORF3a protein QKU30570.1 contains a mutation W131C which was found to be a diseased type and polarity changed from hydrophobic to hydrophilic. This mutation might affect the function of the ORF3a protein.
- QIZ13838.1 possess a mutation L95F which was found to be a diseased type with no polarity change.
- There is a mutation a V55F in QKU29039.1, which was found to be a diseased type with no polarity change. But the mutation may cause an increase in pathogenesis.
- In the protein QKU28847.1, a mutation M260I occurred which was found to be a diseased type with no polarity change and hence functional change of ORF3a can be expected.
- In QLH58085.1, there is a mutation Q185H which was found to be a diseased type with no polarity change and so the structure of ORF3a protein may vary.
- In QKG88539.1, there is a mutation at L108F which was found to be a neutral type with no polarity change. This mutation needs further investigation in order to confirm about its neutrality.
- In QLI50282.1, there is a mutation G18S which was found to be a neutral type, and polarity changed from hydrophobic to hydrophilic. Although this is a neutral mutation but the change in polarity may bear some significance in structural properties.
- In QJU70306.1, there is a mutation at G224C which was found to be a diseased type polarity changed from hydrophobic to hydrophilic. This mutation may change the structure and functions of the protein.
- The ORF3a protein QLI51614.1 contains a mutation V197L which was found to be a diseased type with no polarity change.

In this network flow of mutations, it was also found sequences possessing 3rd level mutations which are described below:

- QLA47776.1: this sequence contains three mutations (Q57H, V55S, A23S), 3rd mutation is the neutral type, and polarity changed from hydrophobic to hydrophilic. Such mutations altogether may affect both structure and function of the protein.
- QJW28665.1: this sequence contains three mutations (Q57H, G224C, L65F), 3rd mutation is the neutral type with no polarity change. The mutation L65F might not affect in virulence property of the SARS-CoV2.

#### Flow of consecutive mutation-III

In this case, network flow (Fig.9) of mutations is devised based on the ORF3a proteins of Indian origin. The sequence QLA10225.1 contains a mutation Q57H as usual. Further five ORF3a proteins are turned up in the network flow in the succeeding time scale of collection of samples. It was found that, all of them possess second mutation along with Q57H.

**Figure 9:**
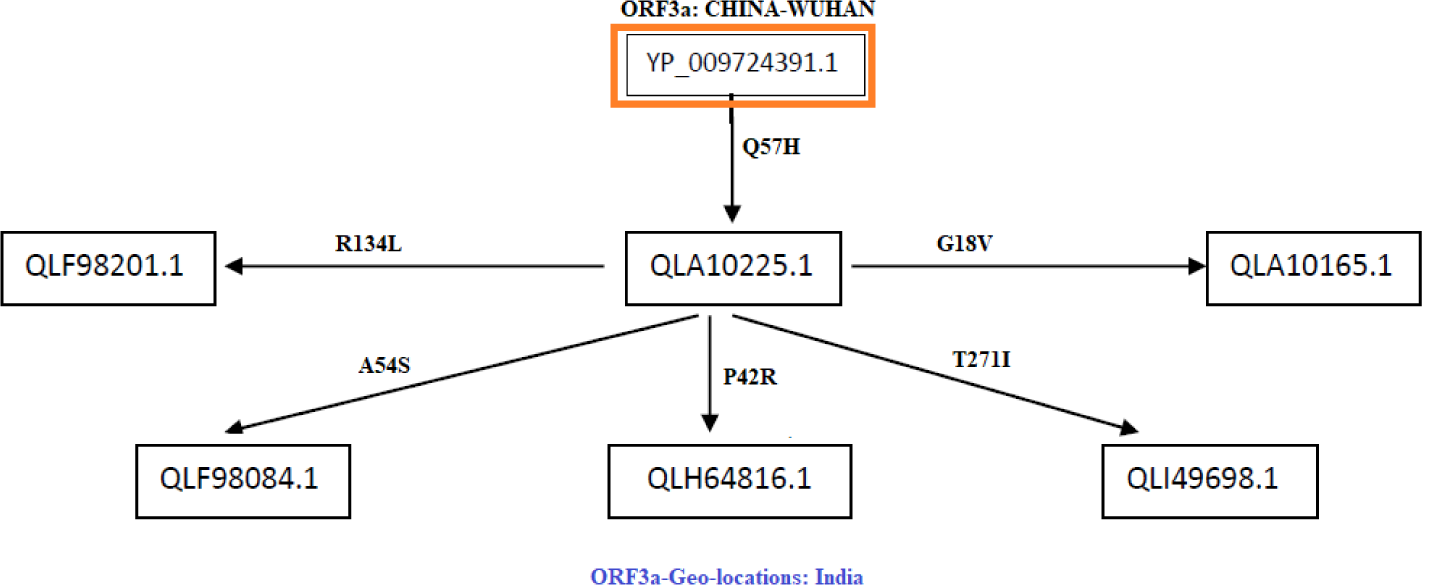
Flow of mutations in ORF3a proteins of Indian origin.

- The mutation R134L in the ORF3a protein QLF98201.1, which was found to be a disease type and there was a polarity change from hydrophilic to hydrophobic. Here the change in mutations may lead to changes in tetramerization properties of the protein.
- The protein QLF98084.1 possesses a mutation at A54S, which was found to be a disease type and the polarity changed from hydrophobic to hydrophilic and hence the structure of the protein is expected to be differed and accordingly the functions of the ORF3a protein would be affected.
- QLH64816.1, there is a mutation at P42R which was found to be a disease type and there was a change in polarity from hydrophobic to hydrophilic and consequently the mutation may contribute to structural changes of the ORF3a protein.
- The protein QLI49698.1 contains the mutation T271I which was found to be a neutral type and there was a change is polarity from hydrophilic to hydrophobic. Although the mutation is predicted to be neutral but the hydrophobicity is changed and hence alternation of functions of the proteins is anyway expected.
- In ORF3a protein QLA10165.1, there is a mutation G18V which was found to be a neutral type of mutation and there is no change in polarity and consequently functions of the proteins would remain same.

#### Flow of consecutive mutation-IV

The sequence QLC46986.1 contains a mutation Q38P which is a disease mutation with the change in polarity from hydrophilic to hydrophobic which might cause a change in functions of the protein. The network flow of mutation id presented in the Fig.10.

**Figure 10:**
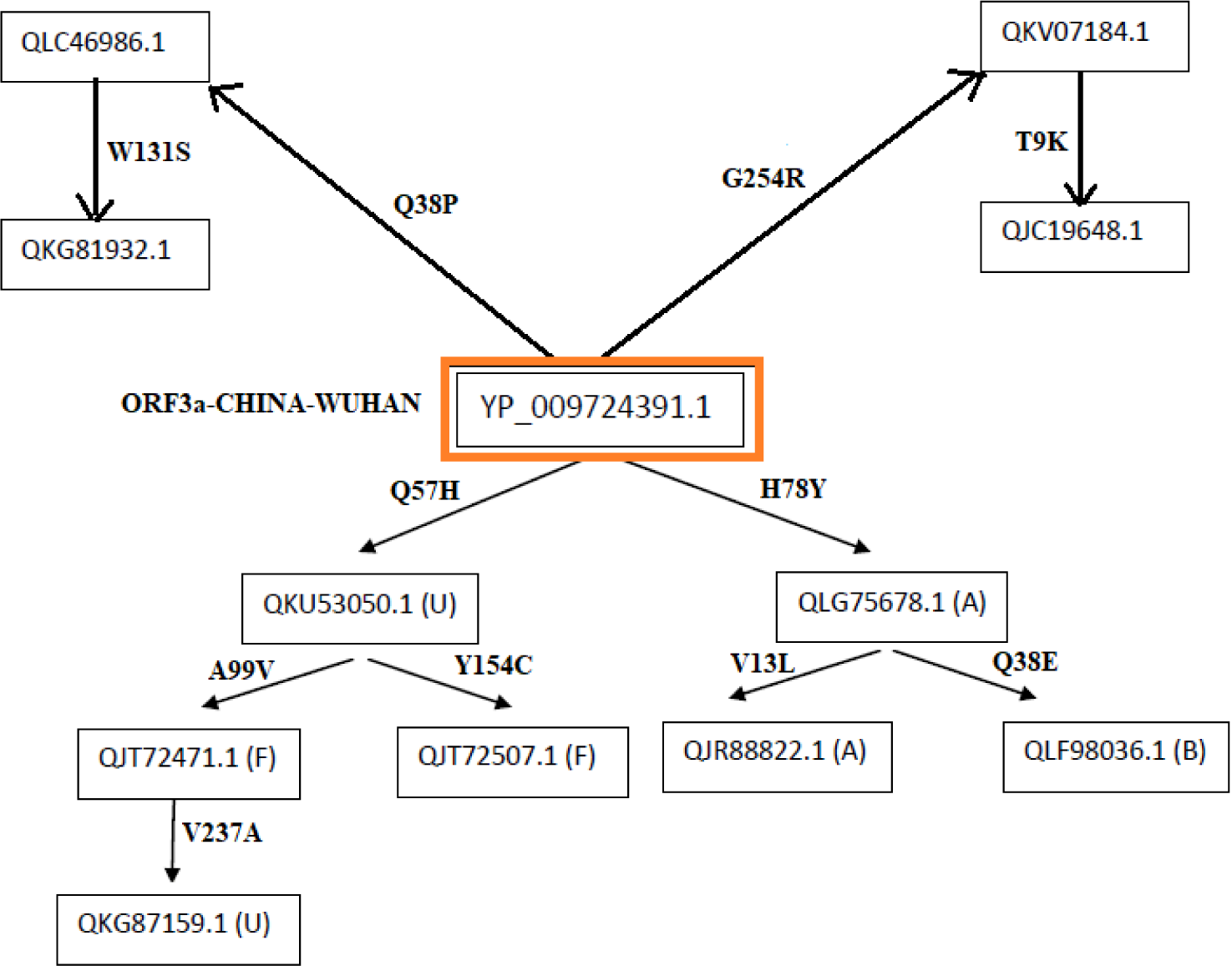
Network flow of mutations of ORF3a proteins considering from various geo-locations.

A second level mutation along with Q38P occurred in QKG81932.1 sequence from W131S which is also a disease type mutation and polarity changed from hydrophobic to hydrophilic and so, it may change the structure of the protein. Also the ORF3a protein QKV07184.1 possesses G254R which changed the polarity from hydrophobic to hydrophilic and caused disease type mutation. On further analysis, the QJC19648.1 sequence was identified to have G254R along with T9K which is a disease mutation with no change in polarity. This is a mutation at the C-terminal region of protein so this mutation may effect the protein-protein interaction.

There is another sequence QKU53050.1(from USA) present in the work flow, which contains the usual mutation Q57H and a France based ORF3a sequence QJT72471.1 possessing a Q57H mutation along with A99V mutation which is a disease type mutation with no change in polarity. QJT72507.1 is another sequence of France origin, in which there is a mutation at Y154C along with Q57H mutation. Also in the QKG87159.1 sequence, another mutation apart from Q57H and A99V at position V237A which is a disease type with no change in polarity.

Another possible traffic of mutation was observed in which an Australian sequence QLG75678.1 had a mutation at 78th position from H to Y, a neutral mutation with no change in polarity which may be a virulence promoting factor. Another Australian sequence QJR88822.1 was identified in which H78Y mutation was observed with V13L which is a disease mutation with no change in polarity. So here we observed that along with a neutral mutation a disease mutation has occurred and it can be assumed that virus first evolved in terms of virulence then enhanced its functional activity. Although there is no change in polarity but it may affect the chemical properties. The sequence QLF98036.1 was another sequence from Bangladesh found to have H78Y mutation in addition to Q38E which is a disease mutation with no change in polarity. here also a disease mutation is observed along with neutral mutation again signifying the evolutionary importance of these mutations.

#### Flow of consecutive mutation-V

The network flow of mutations (Fig.11) with reference sequence of Wuhan’s (ID YP_009724391.1) is formed.

**Figure 11:**
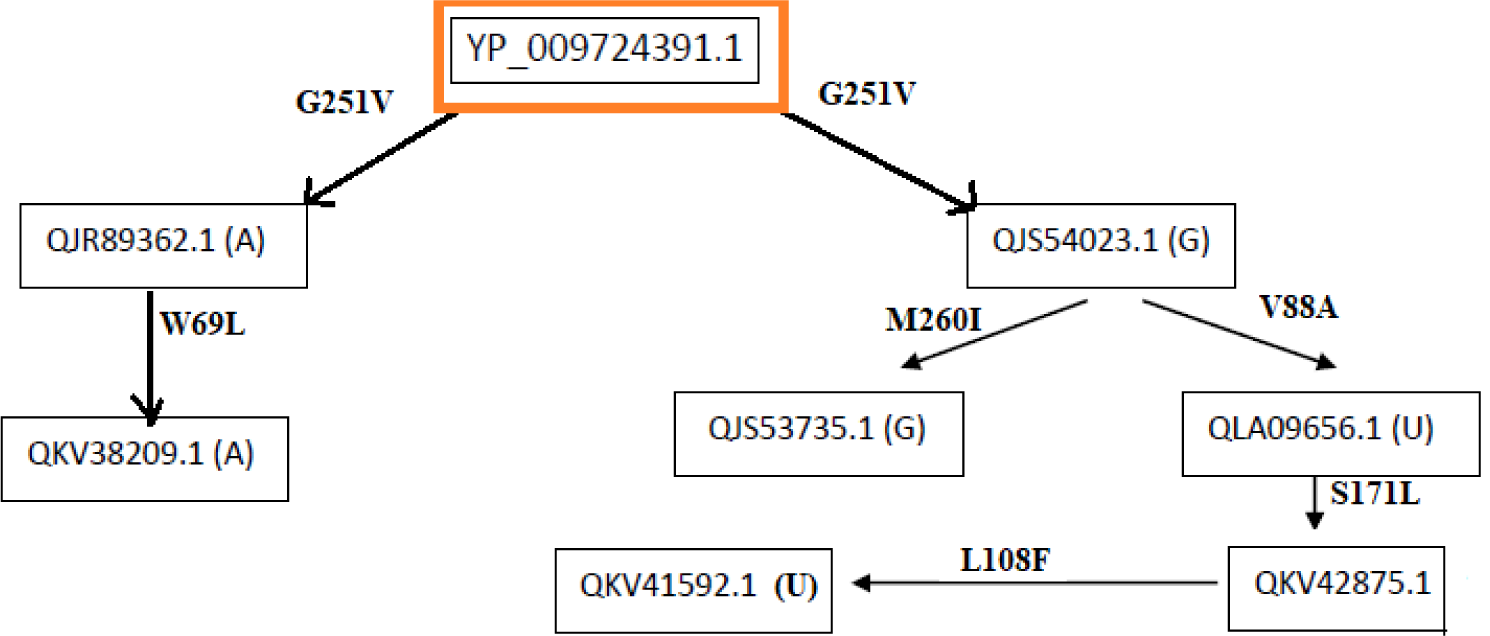
Network flow of mutations of ORF3a proteins considering from various geo-locations. Note: A: Australia, B: Bangaldesh, F: France, G: Greece and U: USA.

The ORF3a protein QJR89362.1 possess a mutation G251V. It was found to be a disease type mutation and here no change in polarity is observed but it may have some significance as it is a disease causing mutation. From this originates another sequence in the flow whose sample collection date is ensuing to the previous one. This sequence (ID QKV38209.1) bears a mutation in W69L which is a disease mutation without any change of polarity that is both W and L are neutral. As this sequence has both the disease mutations, it indicates their functional importance.

In the second case, when the sequence (ID QJS54023.1) of geo-location Greece, is compared with the Wuhan sequence it bore the same mutation G251V. From here it is further divided into bi-flow according to geo-locations and all of them have the G251V mutation along with certain new:

1. The left one bears a sequence (ID QJS53735.1) of geo-location Greece which has a mutation M260I which is a disease type of mutation and has no change in polarity. Here, both the mutations are in the cytosolic domain indicating that these mutations are somehow important for the virus.
2. The right one is for the geo-location USA, which starts with the sequence (ID QLA09656.1) which has a mutation V88A. It is a disease type mutation with no change in polarity. So, it may be advantageous for virus in terms of functionality. Following there is another sequence (ID QKV42875.1) with respect to the time scale, bearing a mutation at S171L. This is a disease type mutation and there is a change in polarity from hydrophilic to hydrophobic. Since the polarity is changing which indicates that there is some effect on ionic and electrostatic interactions that may cause structural changes. Lastly, the sequence QKV41592.1 which bears a mutation at L108F which is a neutral mutation which has no change in polarity. This sequence has all disease mutations although no change in polarity is observed except for one mutation, so it signifies the order of occurrence of mutations allowing the virus to acquire new characteristics important for its survival.

In this study of mutation among many, we recognised five important mutations in the ORF3a proteins. While W131C, T151I, R134L and D155Y forms a network of hydrophobic, polar and electrostatic interactions which are important for the tetramerization process of ORF3a (the functional unit of ORF3a), F230 insertion is responsible for dimerization of ORF3a. We could see that all of the mutations have an effect of decrease in the stability apart from T151I which increases the stability of the protein. To get a better insight, we analysed for these mutations from a structural point of view:

#### Case-I

We collected the available structure of ORF3a (Protein ID: 6XDC) from Protein Data Bank(PDB), (leftmost figure shown in colour grey) in Fig.12

**Figure 12:**
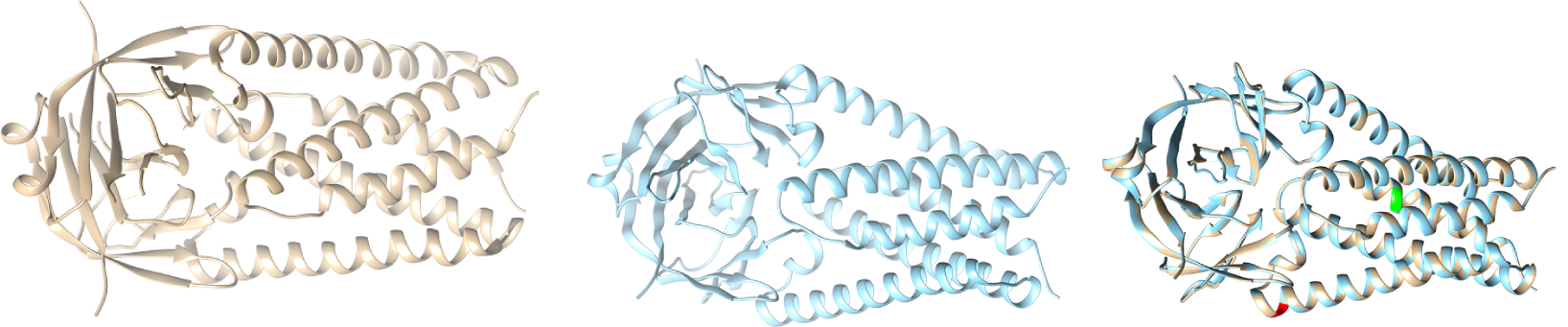
Structures of ORF3a (Reference coloured as grey in left), Structure of mutated ORF3a (coloured with blue in the middle) and Overlayed ORF3a (rightmost image)

Then we took the mutated sequence which contains the mutation W131C and performed homology modelling with the help of a web server called Swiss-model and built the corresponding structure of W131C (middle picture shown in blue) and finally we superimposed the structure of Wuhan (reference structure) with that of the modelled (right most picture) and checked for the corresponding differences with respect to structural change; labelling the mutated portions with colour green(Q57H) and red(W131C).

#### Case-II

In this case, we consider the mutated sequence which possesses the mutation T151Y and performed homology modelling and built the corresponding structure of T151Y (middle picture shown in blue) as shown in Fig.13.

**Figure 13:**
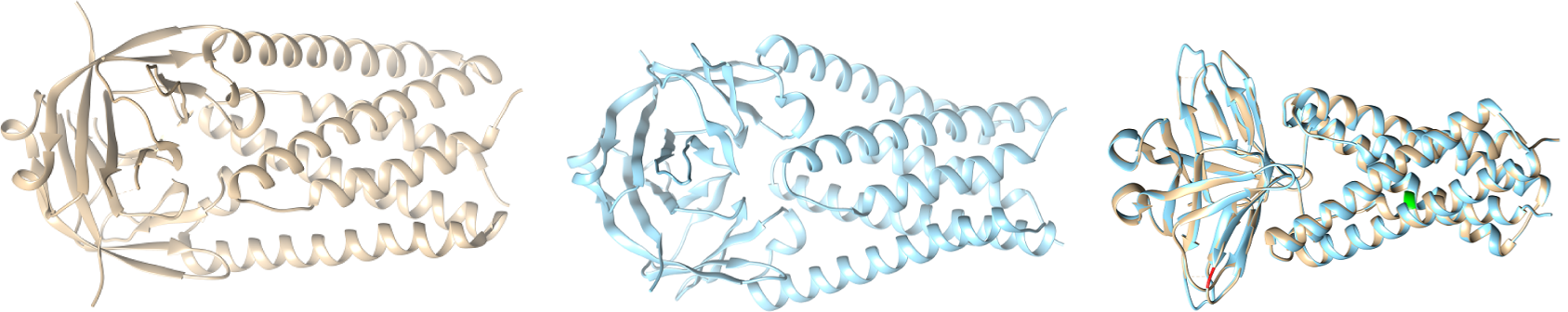
Structures of ORF3a (Reference coloured as grey in left), Structure of mutated ORF3a (coloured with blue in the middle) and Overlayed ORF3a (rightmost image)

Finally we overlayed the structure of Wuhan (reference structure) with that of the modelled (right most picture) and checked for the corresponding differences with respect to structural change; labelling the mutated portions with colour green(Q57H) and red(T151Y).

#### Case-III

With the available structure of ORF3a (Protein ID: 6XDC) from Protein Data Bank(PDB), (leftmost picture shown in colour grey) we took the mutated sequence of R134L and performed homology modelling and built the corresponding structure of R134L (middle picture shown in blue in Fig.14)

**Figure 14:**
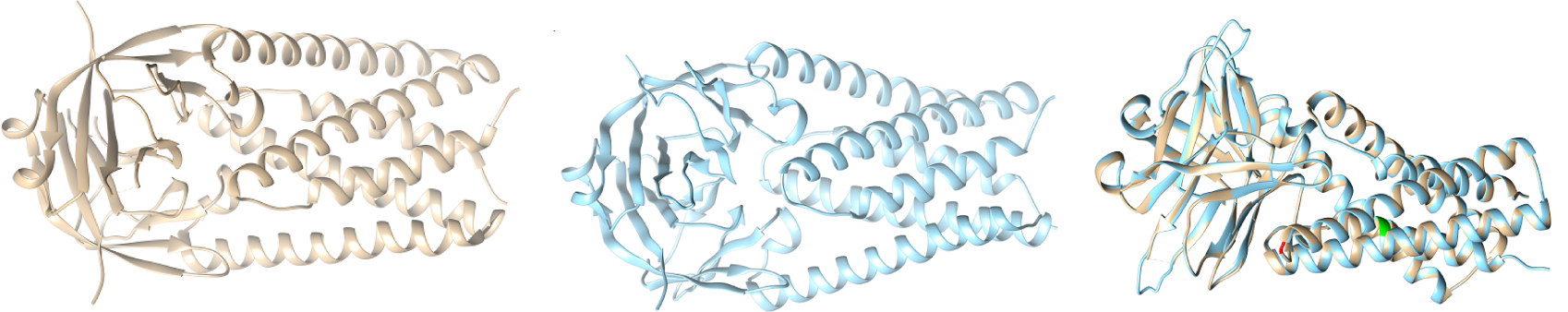
Structures of ORF3a (Reference coloured as grey in left), Structure of mutated ORF3a (coloured with blue in the middle) and Overlayed ORF3a (rightmost image)

Then we overlayed the structure of Wuhan (reference structure) with that of the modelled (right most picture) and checked for the corresponding differences with respect to structural change; labelling the mutated portions with colour green(Q57H) and red(R134L).

#### Case-IV

With the available structure of ORF3a (Protein ID: 6XDC) (leftmost picture shown in colour grey) and then we took the mutated sequence ORF3a considering the mutation D155Y and performed homology modelling and obtained the corresponding structure of D155Y (middle picture shown in blue in the Fig.15).

**Figure 15:**
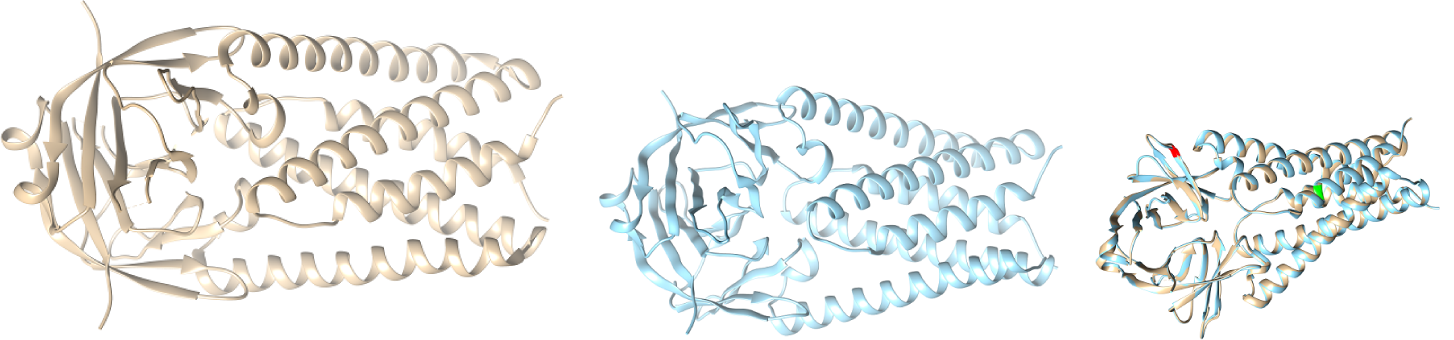
Structures of ORF3a (Reference coloured as grey in left), Structure of mutated ORF3a (coloured with blue in the middle) and Overlayed ORF3a (rightmost image)

We then overlayed the structure of Wuhan (reference structure) with that of the modelled (right most picture) and checked for the corresponding differences with respect to structural change; labelling the mutated portions with colour green(D155Y) and red(D155Y).

#### Case-V

Using the structure of the ORF3a (Protein ID: 6XDC) (leftmost picture shown in colour grey in Fig.16) by homology modelling the structure of the ORF3a protein which contains the insertion mutation F230 (middle picture shown in blue), is constructed.

**Figure 16:**
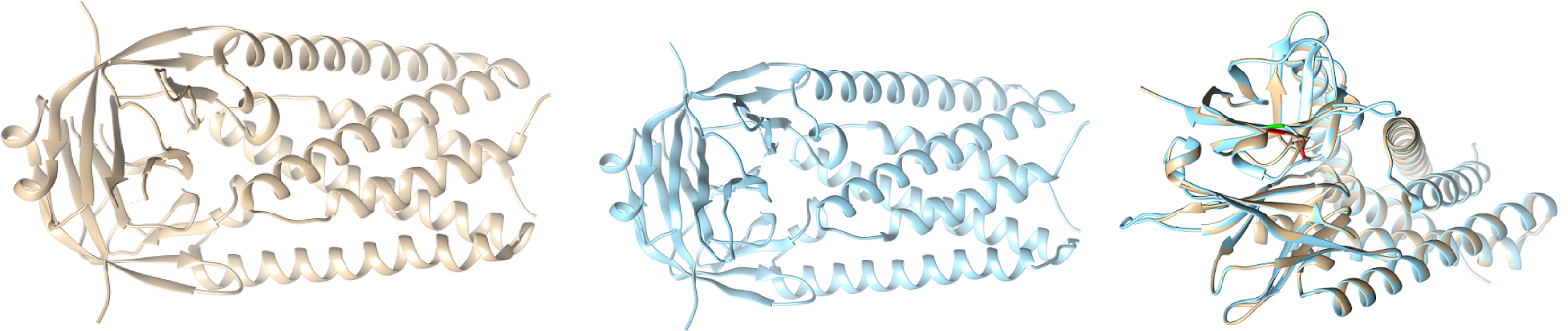
Structures of ORF3a (Reference coloured as grey in left), Structure of mutated ORF3a (coloured with blue in the middle) and Overlayed ORF3a (rightmost image)

Then we overlayed the structure of ORF3a based in Wuhan (reference structure) with that of the modelled (right most picture) and checked for the corresponding differences with respect to structural change; labelling the mutated portions with colour green(difference in structure) and red(inserted amino acid).

We did the above study and no significant change in protein structure was observed, we need a better soft-ware to find the difference between Wuhan sequence and mutated sequences.

### 3.3. Phylogeny and Clustering

We attempted to cluster to cluster each of the 296 ORF3a proteins into twenty disjoint clusters based on the probability distribution of amino acids using K-means clustering technique (Table 12). Note that, the number of clusters (twenty) is chosen optimally by heuristic method in such a manner that the clusters are separated from each other significantly. The frequency probability of each amino acids across all the 296 ORF3a proteins is available as a supplementary file-I. The three truncated ORF3a proteins (detected in Indian patients) are clustered in the cluster 11 as shown in Table 13.

**Table 12:**
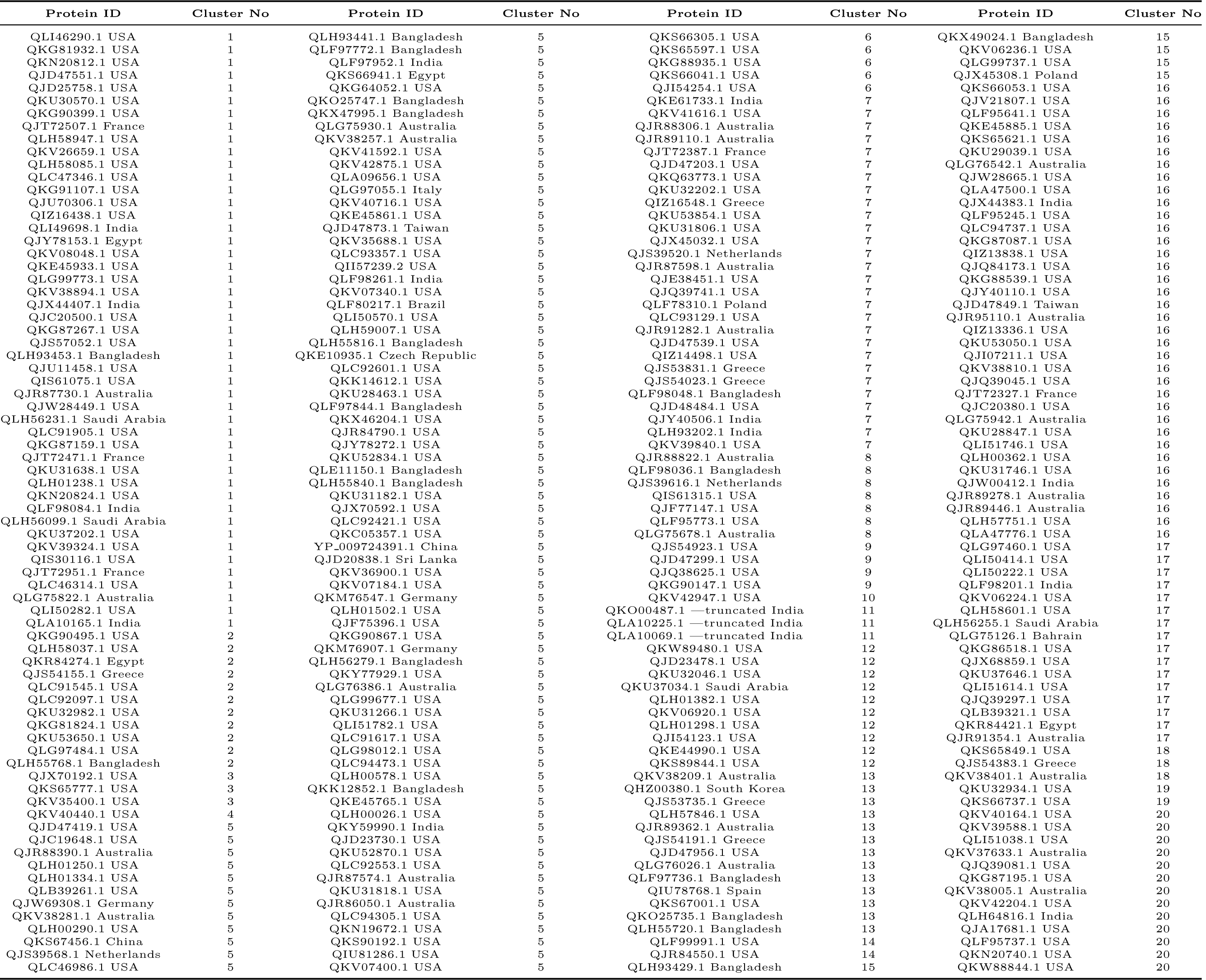
ORF3a proteins and corresponding cluster number based on amino acid distributions.

**Table 13:** Clusters and its frequencies.

| Cluster | Frequency | Cluster | Frequency |
| --- | --- | --- | --- |
| 1 | 47 | 11 | 3 |
| 2 | 11 | 12 | 10 |
| 3 | 3 | 13 | 13 |
| 4 | 1 | 14 | 2 |
| 5 | 86 | 15 | 5 |
| 6 | 5 | 16 | 36 |
| 7 | 28 | 17 | 16 |
| 8 | 7 | 18 | 3 |
| 9 | 4 | 19 | 2 |
| 10 | 1 | 20 | 13 |

The largest cluster 5 contains 53 ORF3a proteins of the USA patients including other 33 from various geo-locations as shown in Table 12. It is found that the ORF3a variants of the USA belong to each of the clusters except the cluster 11 which contained only three truncated proteins belong. This observation confirms the diversity of ORF3a isolates from the USA. It has been seen that the clusters 4, 6, 9 and 10 contains only the ORF3a proteins which are isolated from USA patients. It has been observed that the ORF3a proteins belonging to the clusters 4 and 10 do not possess any mutations and clearly these two ORF3a sequence contain ambiguous amino acids otherwise they would not have appeared as distinct variants.

Based on the hierarchical clustering method, single linkage dendrogram was obtained using the distance matrix of the clusters formed by the K-means clustering method over the 296 ORF3a proteins. This dendogram (Fig.17) depicts the nearness of the clusters which are formed.

**Figure 17:**
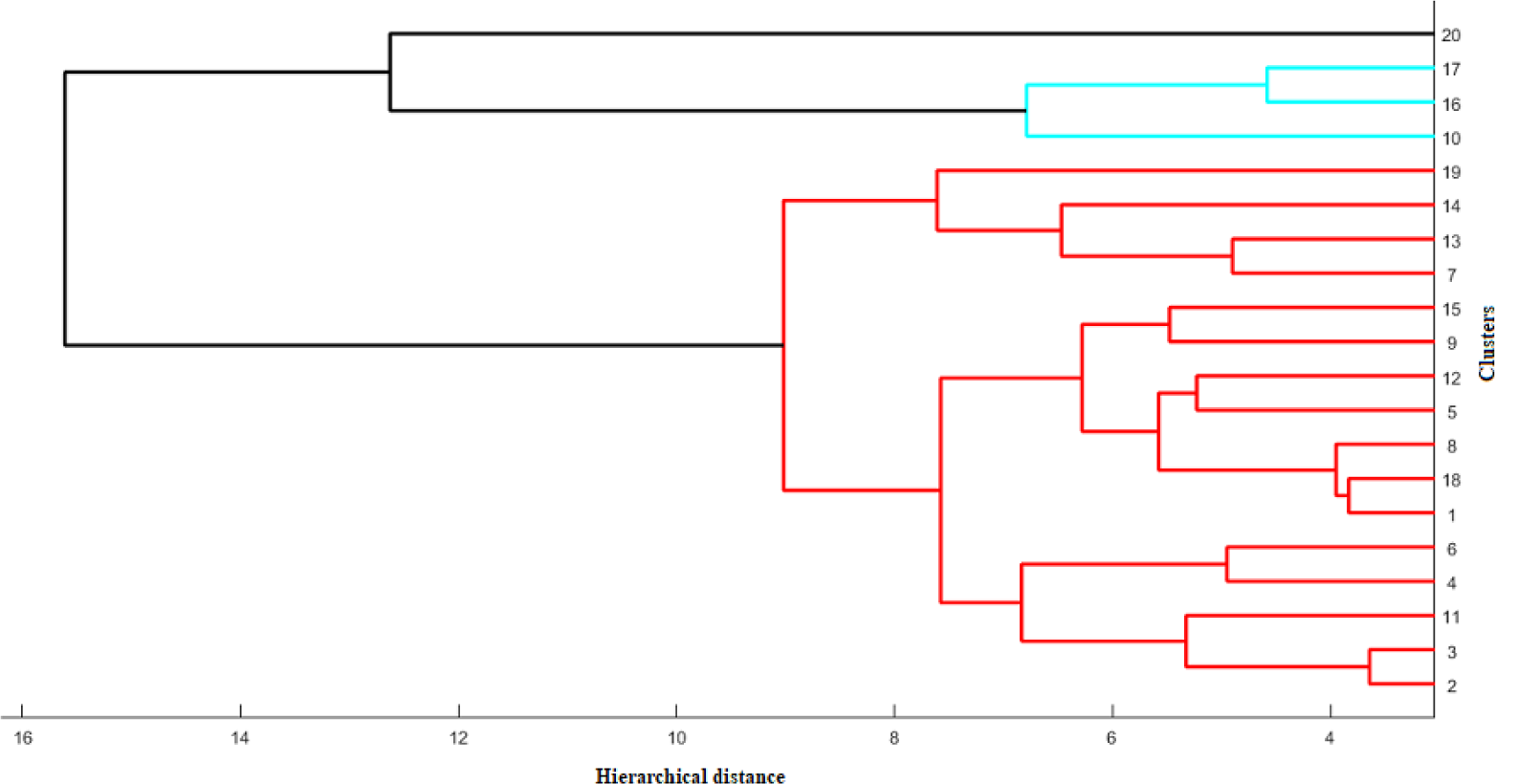
Dendogram of the twenty clusters.

The most nearest pair of clusters are (2, 3), (4, 6), (1, 18), (5, 12), (9, 15), (7, 13) and (16, 17) as observed from the dendogram (Fig.17).

### 3.4. Variability of ORF3a Isolates

The variations among the ORF3a proteins based on the disorderly character of the amino acids over the proteins were determined using Shannon entropy (SE). For each sequence, SE is determined according to the formula stated in the method 2.2.3 and shown in Table 14.

**Table 14:**
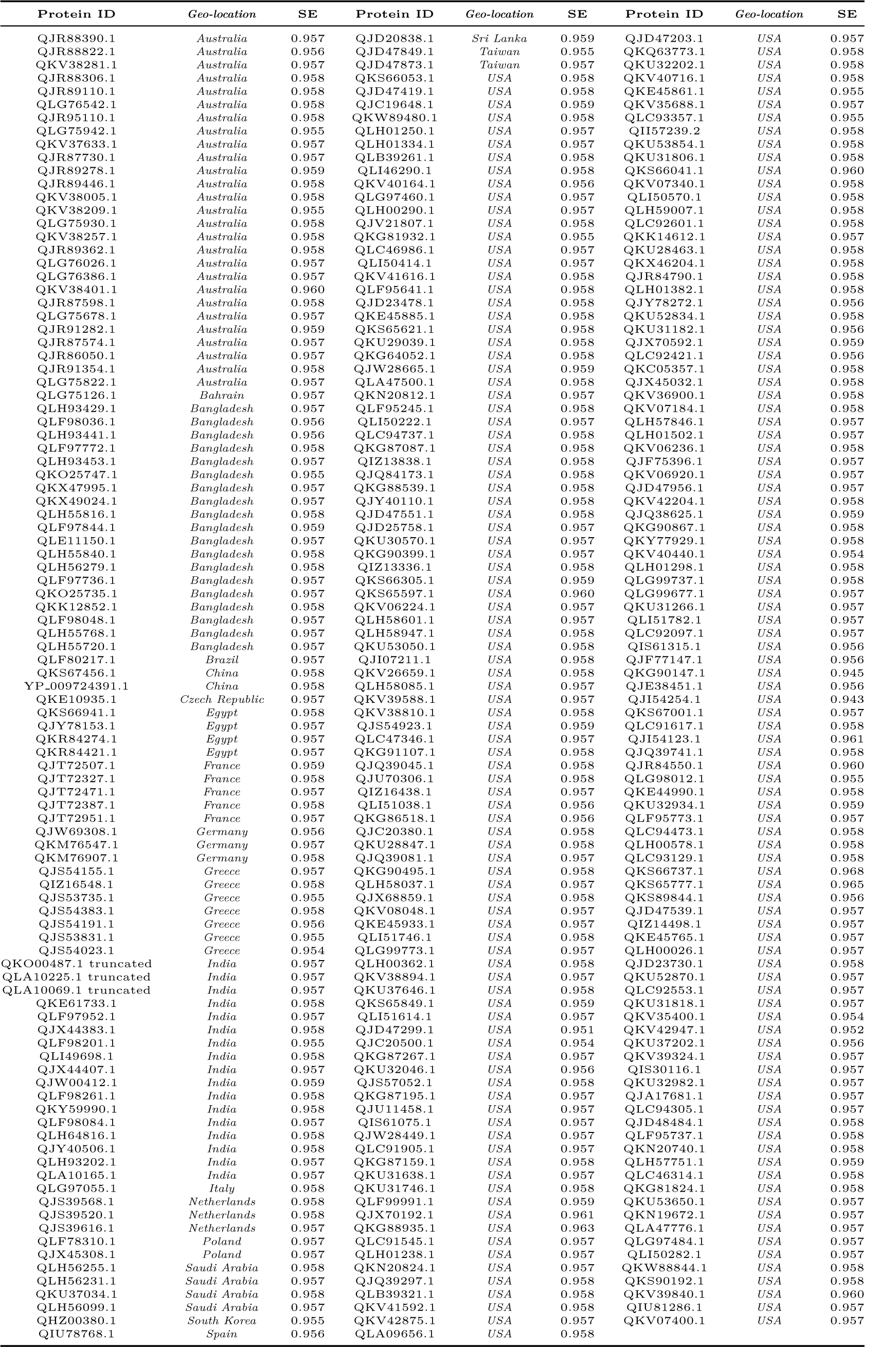
Shannon entropy of amino acid conservations of the 296 ORF3a distinct variants across the world.

The SE of all the ORF3a proteins is bounded by the global minima 0.943 and global maxima 0.968 which are indeed the same as the minima and maxima of the ORF3a proteins which belongs to the USA (Table 15). Clearly, the amount of disorderliness of the amino acids over the ORF3a proteins is extremely high.

**Table 15:** Maxima and minima of SEs across geo-locations.

| *Geo-location* | *Min* | *Max* | *Range* |
| --- | --- | --- | --- |
| Australia | 0.955 | 0.96 | 0.005 |
| India | 0.955 | 0.959 | 0.004 |
| **USA** | **0.943** | **0.968** | **0.025** |
| Bangladesh | 0.955 | 0.959 | 0.004 |

The range of SE of the ORF3a proteins of SARS-CoV2 collected from USA is comparatively more than others and it ensures the wide variety of distinct ORF3a in USA patients. The SEs of 296 ORF3a proteins are plotted (Blue line) in the Fig.18. We found various non-smooth peaks and those are clearly the SEs of the ORF3a proteins of the USA patients and that is reconfirmed in the SE plot (Red line) of the ORF3a proteins of the USA.

**Figure 18:**
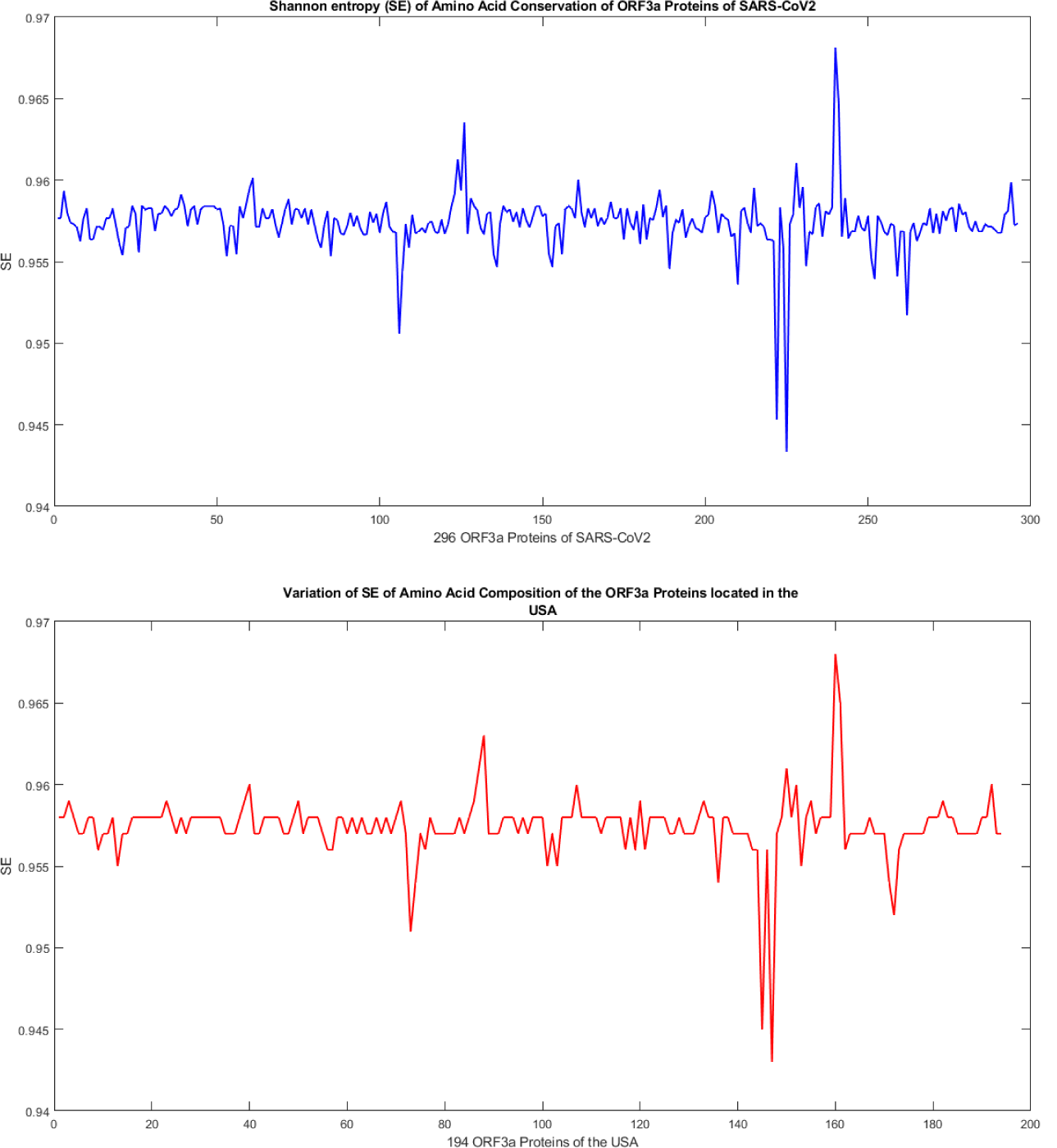
SE of amino acid compositions of ORF3a proteins.

## 4. Discussions

A total of 175 distinct mutations across the distinct 256 ORF3a proteins of SARS-CoV2 are detected and further analyzed. Among all the mutations, 32 mutations were already reported [27, 23]. It was reported that in SARS-CoV, there exists an intensive interchain disulphide bonds with that of the spike protein with the help of the cysteine residues present in ORF3a protein. SARS-CoV2 ORF3a, contains a similar functional region (Domain III: C133) which is found to be conserved, as we did not find any mutation in this region. So, it can be assumed that these cysteine domains perform a similar function as in SARS-CoV and is functionally important for virulence. In SARS-CoV, it was reported that tetramerization of the ORF3a protein is an important step for the ion channel formation which further increased the infectivity of the virus. From this study we found mutations W131C, T151I, R134L and D155Y which may facilitate the tetramerization process in SARS-CoV2 and thereby assisting the ion channel formation and favouring the virus with its infectivity. Similar to that of SARS-CoV, it is also responsible for apoptosis mediated by TRAF-3 (Domain III). We found two mutations in this region Q38E and Q38P which may enhance the effect of apoptosis but further studies are required. Caveolin-binding domain is responsible for viral uptake of the host cell and its translocation to various endomembrane organelle. We have also isolated mutations in this zone (C148Y and A143S) which may enhance the viral uptake by the host, thereby increasing the infectivity rate. However, it is noteworthy that in *Y XXϕ* motif domain, no mutation is observed so far and consequently this domain is conserved. In seven ORF3a variants from the USA, two mutations are found in SGD domain (S171L & G172C), however the function of this SGD domain is unknown.

We characterized the mutations into three types: Neutral, Disease and Mixed. Among these three mutations we found that disease mutations are highly prevalent with a percentage of 66% in the geo-loaction of USA, indicating disease-causing character of the virus getting intensified and thus posing threat to mankind. Simultaneously, we have the mixed type mutation occurring with a rate of 79% in the geo-location of USA. Mixed type had both disease and neutral occurring together. Although, neutral mutations are there in mixed type but frequency of disease mutation is high, again pointing towards the viral advantage over host. In France although the infectivity rate was very high, but disease (2.9%) mutation rate was low compared to USA; where we find the maximum variety of mutation as shown with Shannon entropy in this study. So, we can suggest that the possible wide variety of mutations in USA is due to the high rate of travel within USA and from outside USA, while in France there might be within-country transmission which resulted in less frequent mutations. We also checked the mortality rate of USA (3.3%), France (13.4%) and India (2.1%) and from the results we found that France has the highest mortality rate than USA followed by India. So, consequently we can draw a conclusion that since France has only disease type mutation unlike that of USA and India in which all three type of mutations are present. This may prove that the presence of only disease type of mutation in a sequence may pose more danger to mankind than a sequence containing either mixed type or neutral type of mutations. Next, we analysed consecutive mutations within a protein sequence on the basis of chronological order of the time-line of sample collection from COVID-19 infected patients.

We further went on to analyse the mutations responsible for tetramerization and dimerization with respect to structure and found that there were no significant structural changes observed by homology modelling method. So, other method should be used to detect the effect of mutations on the 3D structure of the protein and results need to be experimentally validated. Finally, twenty clusters are formed from 296 distinct variants of ORF3a of SARS-CoV2 based on the amino acid compositions of the proteins. It also shows wide variety of compositions of ORF3a variants in the USA which is further quantitatively supported by the SE. This study of comprehensive 175 novel mutations would help in understanding the pathogenetic contribution of the ORF3a proteins. This understanding is an important aspect in devising vaccine for COVID-19.

## Author Contributions

SH conceived the problem. DA, SG and SH examined the mutations. All the authors analysed the data and result.

SH wrote the initial draft which was checked and edited by all other authors to generate the final version.

## Acknowledgement

^ϒ^Ms. Diksha Attrish and ^†^Ms. Shinjini Ghosh are Interns under the supervision of Dr. Sk. Sarif Hassan through Virtual Internship with Science Leader (VISL) Programme, 2020. Authors thank to the Virtual Internship with Science Leader (VISL) program for their supports.

